# LncRNA *RUS* shapes the gene expression program towards neurogenesis

**DOI:** 10.1101/2022.02.17.480853

**Authors:** Marius F. Schneider, Veronika Müller, Stephan A. Müller, Stefan F. Lichtenthaler, Peter B. Becker, Johanna C. Scheuermann

## Abstract

The evolution of brain complexity correlates with an increased expression of long, non-coding (lnc) RNAs in neural tissues. Although prominent examples illustrate the potential of lncRNAs to scaffold and target epigenetic regulators to chromatin loci, only few cases have been described to function during brain development. We present a first functional characterization of the lncRNA *LINC01322*, which we term RUS for ‘<u>R</u>NA <u>u</u>pstream of <u>S</u>litrk3’. The *RUS* gene is well conserved in mammals by sequence and synteny next to the neurodevelopmental gene Slitrk3. *RUS* is exclusively expressed in neural cells and its expression increases along with neuronal markers during neuronal differentiation of mouse embryonic cortical neural stem cells. Depletion of *RUS* locks neuronal precursors in an intermediate state towards neuronal differentiation resulting in arrested cell cycle and increased apoptosis. *RUS* associates with chromatin in the vicinity of genes involved in neurogenesis, most of which change their expression upon *RUS* depletion. The identification of a range of epigenetic regulators as specific *RUS* interactors suggests that the lncRNA may mediate gene activation and repression in a highly context-dependent manner.

## Introduction

Most parts of a higher eukaryotic genome are transcribed at times and in certain cells, but only a minority of the resulting RNAs are protein-coding. While many of these non-coding transcripts are immediately degraded, others are processed into small RNAs that form an intricate network regulating gene expression in a co- and post-transcriptional manner. In addition, mammalian genomes encode thousands of stable RNAs longer than 200 nucleotides, often capped and polyadenylated, but without any obvious coding potential [long, noncoding (lnc) RNAs] (Engreitz *et al*, 2016; Quinn & Chang, 2016; Rutenberg-Schoenberg *et al*, 2016; Kopp & Mendell, 2018). The functions of most lncRNAs discovered in large-scale sequencing projects remain to be explored. ‘Guilt-by-association’ strategies correlate their presence and expression levels with certain cellular states, including disease conditions. Increasingly, interference strategies reveal critical roles for lncRNAs in cellular fates and states (Statello *et al*, 2021; Rinn & Chang, 2020; Lin *et al*, 2014).

Apparently, lncRNAs arise by pervasive transcription of the genome and evolve fast. Conceivably, their structural flexibility makes them an ideal substrate for ‘constructive neural evolution’ and predisposes them for a function in chromatin regulation (Palazzo & Koonin, 2020; Rinn & Chang, 2020). Indeed, more than 60% of annotated lncRNAs in human cells are chromatin-enriched (Rinn & Chang, 2012).

In the chromatin context, lncRNAs often combine two functions: scaffolding and targeting. The intrinsic ability of lncRNAs to mediate positional targeting in the genome qualifies them to impose allele-specific epigenetic regulation, such as genome imprinting, X chromosome inactivation or rDNA regulation (Yao *et al*, 2019; Rinn & Chang, 2020; Statello *et al*, 2021). Their actions may be locally restricted close to their site of transcription in *cis*, or in *trans* via sequence-specific hybridization with DNA or RNA. Thus, they may guide powerful ‘epigenetic’ regulators (enzymes that modify histones or DNA) to specific loci in chromatin, or participate in nuclear condensates (Rutenberg-Schoenberg *et al*, 2016; Kopp & Mendell, 2018; Statello *et al*, 2021; Engreitz *et al*, 2016). Prominent examples of lncRNAs recruiting regulators that define epigenetic chromatin states, include *XIST*, *HOTAIR* and *ANRIL* that bind polycomb complexes (PRC) to silence chromosomal regions, while others such as HOTTIP or certain enhancer RNAs are known to recruit activating histone acetyltransferase or methylase complexes (Werner & Ruthenburg, 2015; Quinn & Chang, 2016).

The fraction of lncRNAs that are expressed in a tissue-specific manner exceeds that of cell type-specific protein-coding genes (Djebali *et al*, 2012). A particular rich compendium of lncRNAs is expressed in the mammalian brain (estimated 40% of known lncRNAs) (Mercer *et al*, 2010; Hezroni *et al*, 2019; Briggs *et al*, 2015), and a strong correlation between the number of expressed lncRNAs and mammalian brain size was reported (Clark & Blackshaw, 2017). Brain-specific lncRNAs tend to be more evolutionary conserved between orthologues than lncRNAs expressed in other tissues and their genes often reside next to protein-coding genes involved in neuronal development or brain function processes (Ponjavic *et al*, 2009). Indeed, lncRNAs are drivers of key neurodevelopmental processes such as neuroectodermal lineage commitment, proliferation of neural precursor cells, specification of the precursor cells, and the differentiation of precursor cells into neurons (neurogenesis) or other neural cell types (gliogenesis) (Briggs *et al*, 2015; Zimmer-Bensch, 2019).

Diverse mechanisms have been documented. For example, lncRNA *TUNA* (*megamind*) is involved in neural differentiation of mouse embryonic stem cells (mESCs) (Lin *et al*, 2014). The finding that depletion of *TUNA* also compromised ESC proliferation and maintenance of pluripotency illustrates the power of lncRNA to control gene networks in diverse ways, depending on the nature of protein effectors and the timing and context of their lncRNA interactions (Lin *et al*, 2014). The lncRNA *RMST* promotes neuronal differentiation by recruiting the transcription factor Sox2 to promoters of neurogenic genes (Ng *et al*, 2013). The lncRNA *Pinky* is expressed in the neural lineage, where it helps to maintain the proliferation of a transit-amplifying cell population, thereby restraining neurogenesis. This regulation takes place at the level of transcipt splicing, illustrating the versatility of nuclear lncRNAs (Ramos *et al*, 2015). Other mechanisms involve the control of miRNA availability and function, as has been shown for the primate-specific lncND during neurodevelopment (Rani *et al*, 2016).

Only a small fraction of lncRNAs involved in neurodevelopment and brain function has been studied in detail. We here describe a novel lncRNA involved in neurogenesis, which we term *RUS* (for ‘<u>R</u>NA <u>u</u>pstream of <u>S</u>litrk3’). The *RUS* gene resides at a syntenic position in mouse and human genomes upstream of the Slitrk3 gene, which encodes a transmembrane protein involved in suppressing neurite outgrowth. *RUS* is expressed in neural tissues only and its expression increases during the differentiation of neural stem cells into neurons. *RUS* is a nuclear lncRNA that interacts with chromatin in the vicinity of genes involved in neurogenesis. Depletion of RUS results in massive alterations in the gene expression program of neuronal progenitor cells, trapping them in an intermediate state during differentiation and eventually leading to proliferation arrest. Proteomic identification of *RUS*- interacting proteins suggests multiple mechanisms of *RUS*-mediated epigenetic gene regulation.

## Results

### Identification of the neuronal-specific lncRNA RUS

To identify novel, functionally relevant lncRNAs in the context of neurogenesis, we searched for transcripts lacking obvious coding potential meeting the following criteria. They should 1) only be expressed in neural tissues, 2) be dynamically regulated during the differentiation of neural precursor cells and 3) be conserved between mouse and humans (Fig 1A). We examined the expression of 553 candidate lncRNAs in human ESC-derived neural progenitor cells: neuroepithelial cells (NE), early, mid and late radial glia cells (ERG, MRG, LRG, respectively) before and after differentiation (Ziller *et al*, 2015). Of these, 10 transcripts decrease and 29 increase during the differentiation of the four cell types (Fig 1B). Among them, we identified *LINC01322* as an interesting candidate, as it was absent in NE, ERG and MRG, but expressed in all differentiated cell types. Intriguingly, *LINC01322* was also expressed in undifferentiated LRG.

**Figure 1.**
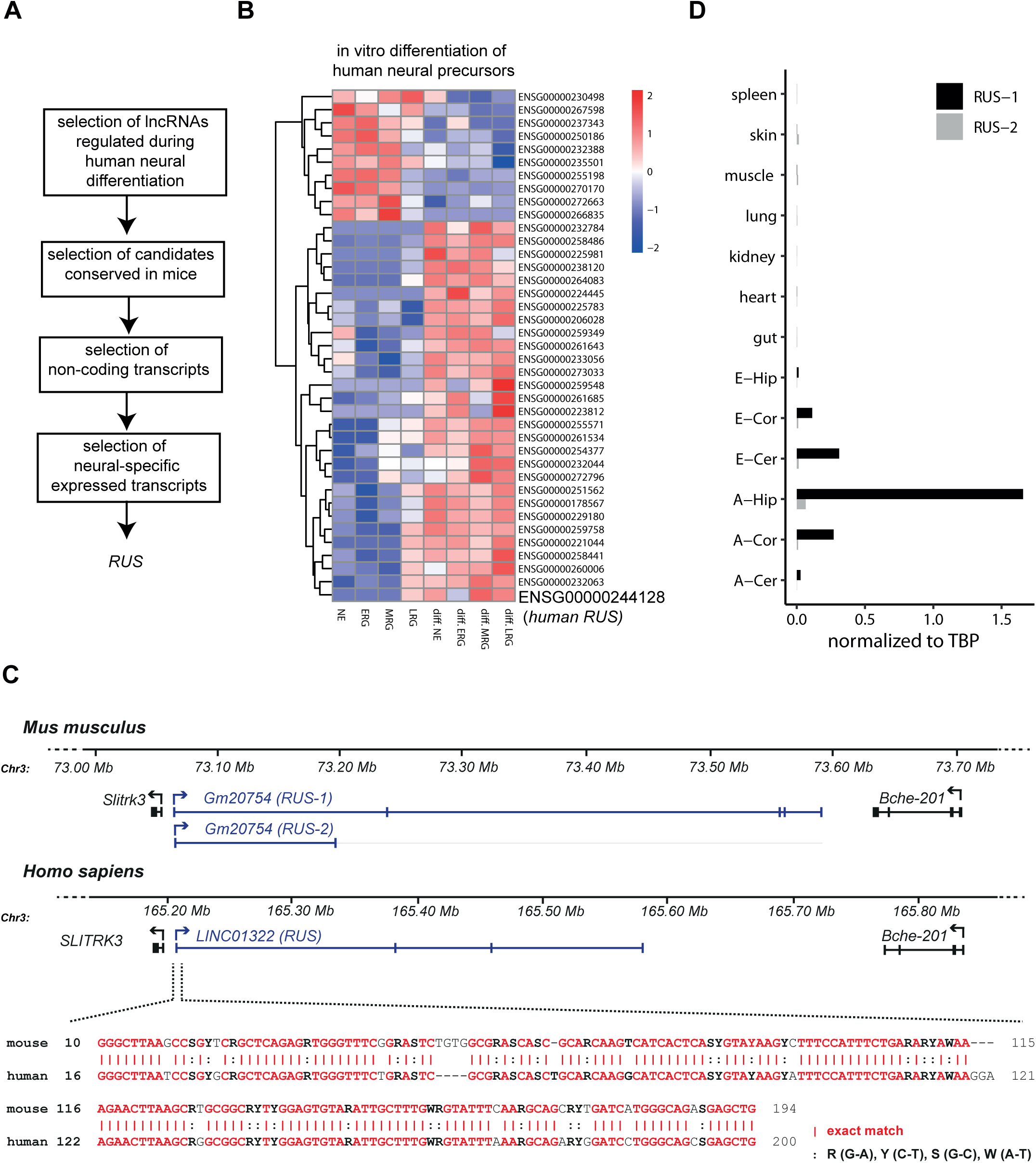
RUS is a novel, conserved lncRNA involved in neurogenesis. **A.** Workflow illustrating the criteria and selection principle applied to candidate lncRNAs (Ziller et al., 2015), which led to the selection of *RUS* as subject of this study. **B.** Heatmap of significantly changed lncRNAs expressed in human ESC-derived NE, ERG, MRG and LRG before and after differentiation (two-sided t-test). Data of (Ziller et. al. 2015) were analysed. **C.** Conservation of the *RUS* gene between mouse and human genomes by synteny (top) and by sequence of exon 1 (bottom). Note that the RUS gene resides just upstream of the *Slitrk3* gene in either case. For mice two *RUS* isoforms are indicated. **D.** Expression of murine *RUS-1* and *RUS-2* isoforms (see panel C) in different embryonic (E-) and adult (A-) tissues: cortex (Cor), cerebellum (Cer), hippocampus (Hip), gut, heart, kidney, liver, lung, muscle, skin, spleen, analyzed by quantitative RT-PCR. The values were normalized to expression constitutive TBP mRNA (arbitrary units).

LncRNA genes relevant to neurogenesis are often located next to neurodevelopmental protein-coding genes (Ponjavic *et al*, 2009). In line with this observation, the gene for *LINC01322* localizes upstream of the gene encoding the transmembrane protein Slitrk3, which regulates neurite outgrowth (Aruga *et al*, 2003) (Fig 1C). In the following, we refer to *LINC01322* as *RUS* (*RNA upstream to Slitrk3*). The location of the *RUS* gene is well conserved by synteny in mice and humans between the *Slitrk3* and *Bche-201* genes (Fig 1C).

The murine *RUS* transcript, *Gm20754,* has two annotated isoforms. Two and five exons are annotated for isoforms 1 and 2, respectively. Both isoforms share the 232 bp exon 1, which is 75% similar to the orthologous counterpart in humans (Fig 1C). The sequence of *mRUS* exon 2 (114 bp) is conserved to 92%, but not part of the predominant human transcript. I*n silico* open reading frame (ORF) predictions revealed that the largest ORF encodes a theoretical polypeptide of 80 amino acids (aa). Since the corresponding peptides are not listed in published mass spec data (PeptideAtlas), we assume that *RUS* functions as a lncRNA.

Quantitative RT-PCR analysis of the two isoforms in different mouse adult and embryonic tissues revealed that *RUS* annotated isoform 1 is the dominant form (Fig 1D). *RUS* expression is restricted to neural tissues, with highest expression in the adult hippocampus. Continuing with isoform 1, we performed 3’-RACE experiments to obtain the annotated 3’ end (Fig EV1A). However, amplification of *RUS* with primers targeting the annotated 5’ and 3’ ends yielded two PCR bands of 1.3 kbp and 0.9 kbp. Sequencing the more abundant 0.9 kbp PCR band revealed that it lacked exon 4. (Fig EV1B).

### RUS depletion leads to reduced neuronal differentiation, proliferation arrest and increased apoptosis

To monitor the expression of *RUS* during murine neurogenesis, we differentiated embryonic cortical neural stem cells (NSC) into immature neurons *in vitro* (Kilpatrick & Bartlett, 1993; Azari *et al*, 2011; Mukhtar *et al*, 2020). Differentiating NSC were maintained proliferative by mitogen (bFGF) for the first 4 days. On day 5, bFGF was withdrawn to induce neurogenesis (Fig EV2A). During a time course of 9 days the expected changes in molecular marker expression were detected via immunostaining and quantitative RT-PCR (RT-qPCR) analyses. The high expression of the NSC marker nestin decreased, with a concomitant increase in RGC markers Gfap, Glast and GluL (Fig 2A, EV2A,B), as observed elsewhere (Imura *et al*, 2003; Mamber *et al*, 2012). Upon bFGF withdrawal, the culture acquired neuronal features with high expression of the neuronal markers Map2, Dcx, β-tubulin III and Mapt (Fig 2A, EV2A-B). The expression level of *RUS* continually increased along with the neuronal markers, reaching robust expression on day 5 of the differentiation process (Fig 2A, EV2B).

**Figure 2.**
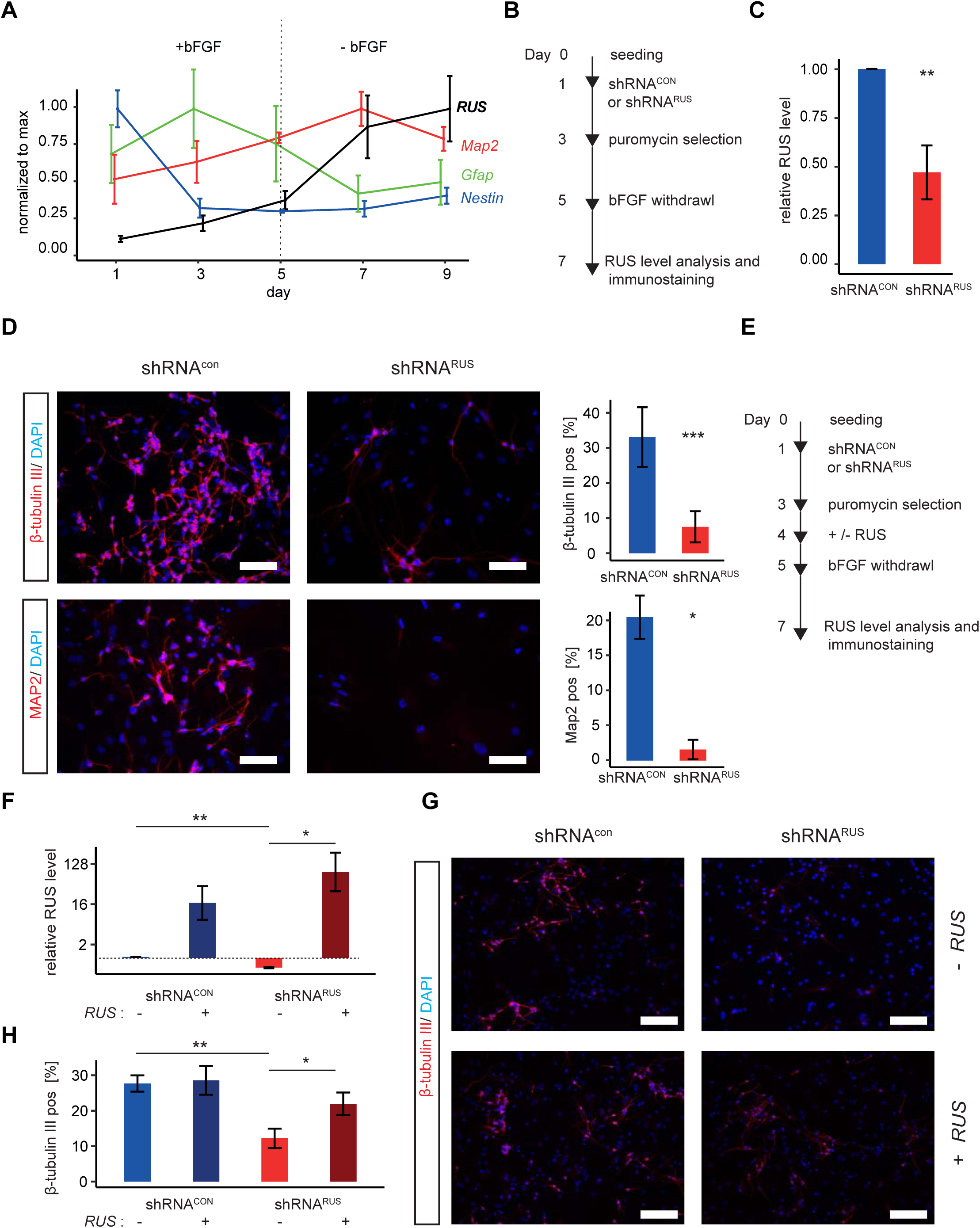
*RUS* is involved in neuronal development. **A.** Quantitative RT-PCR analysis of expression of *RUS*, *Map2*, *Gfap* and *Nestin* transcripts as indicated, during a 9-day time course of embryonic cortical NSC differentiation. Values were normalized to the maximal expression of each RNA during the time course. Error bars show the standard error of the mean of 3 independent experiments. FGF: Fibroblast growth factor. **B.** Experimental strategy to deplete *RUS* in differentiating NSC by expressing shRNAs upon lentiviral transduction. **C.** *RUS* levels determined by RT-qPCR in *RUS* knockdown cells (red, expressing shRNA*^RUS^*) compared to control cells (blue, expressing a scrambled shRNA*^CON^*). Error bars show the standard deviation of the mean of four individual experiments. **D.** Left: Immunofluorescence visualization of β-tubulin III (upper panel) and Map2 (lower panel) in control (shRNA*^CON^*) and knockdown (shRNA*^RUS^*) cells using specific antibodies (magenta). Nuclei were stained with DAPI (4′,6-diamidin-2-phenylindol, blue). Scale bar = 25 µm. Right: Quantification of percentage of immune-positive cells by ImageJ. The bar diagrams show the percentage of positive cells. Error bars show the standard deviation of four independent experiments. **E.** Experimental strategy to rescue the *RUS*-depletion phenotype in differentiating NSC by lentiviral overexpression of *RUS*. **F.** *RUS* levels were determined by RT-qPCR in control (shRNA*^CON^*) and knockdown (shRNA*^RUS^*) cells. Where indicated (+), *RUS* was overexpressed from a CMV promoter. Error bars show the standard deviation of four independent experiments. The dashed line highlights the level of *RUS* in (shRNA*^RUS^*) cells **G.** β-tubulin III immunostaining in control (shRNA*^CON^*) and knockdown (shRNA*^RUS^*) cells as a function of *RUS* overexpression. Nuclei are stained with DAPI. Scalebar = 50 µm) **H.** Quantification of β-tubulin-III immunostaining of cultures as in G. Error bars show the standard deviation of four independent experiments (* p < 0.05, ** p < 0.01, *** p < 0.005).

To explore a potential involvement of *RUS* during neuronal differentiation, we depleted *RUS* by RNA interference, expressing a *RUS*-targeting shRNA (shRNA*^RUS^)* upon lentiviral transduction into differentiating NSC [Fig 2B, Table EV1, (Moffat, et al., 200*6)]. The shRNA^RUS^*was selected to have no predicted off-targets, while significantly reducing RUS levels. Upon expression of shRNA*^RUS^, RUS* levels were typically reduced by approximately 50% compared to control cells expressing a scrambled control shRNA*^CON^* (Fig 2C). Remarkably, upon *RUS* depletion the number of cells expressing the neuron-specific β- tubulin III or the dendritic marker Map2 were reduced to 37% and 8%, respectively (Fig 2D).

The specificity of the knockdown was assessed by a rescue experiment. *RUS*-depleted and control cells were transduced with lentiviruses expressing *RUS* driven by the strong CMV promoter (Fig 2E). RT-qPCR revealed that *RUS* was increased roughly 20-fold compared to endogenous, wildtype levels (Fig 2F). Immunostaining of the cells for β-tubulin III served as a proxy for neurogenesis (Fig 2G). *RUS* expression in cultures that had been depleted of endogenous RUS largely restored the number of β-tubulin III-positive cells but did not further increase this value in the presence of endogenous *RUS* (Figure 2H).

*RUS* depletion led to reduced cell numbers in culture, which may be a consequence of reduced cell proliferation or increased apoptosis. To explore whether this cell loss was due to reduced cell proliferation, we supplemented differentiating NSC cultures with BrdU and monitored its incorporation by immunostaining as a measure of replication (Figure EV2C,D). *RUS* depletion reduced the number of BrdU-positive, proliferating cells by 93.7% (Figure EV2D). We also probed for apoptosis. We replaced the puromycin resistance gene in the shRNA vector by a GFP gene to visualize knockdown cells while avoiding cell death due to puromycin selection (Figure EV2C). Immunostaining for cleaved caspase 3 in GFP-positive cells revealed a 9-fold increase of apoptosis in shRNA*^RUS^*-expressing cells compared to a very low level in control cultures (Figure EV2E). We conclude that the depletion of RUS in differentiating NSCs inhibits cell proliferation and induces apoptosis.

### Depletion of RUS locks neural progenitor cells in their differentiation stage

For an in-depth characterization of the shRNA*^RUS^*knockdown phenotype in differentiating NSC we monitored transcriptional changes by RNA-seq analysis. We established the transcriptome at days 5 and 7 after seeding, when endogenous RUS expression is drastically increased, in cells either treated with shRNA*^RUS^* or shRNA*^CON^* (Fig EV3A, Table EV2). RNA interference by shRNA*^RUS^* reduced *RUS* levels to roughly 50%, as before (Fig EV3B). Despite this incomplete depletion, the principal component analysis (PCA) of four replicates clearly separated shRNA*^CON^* and shRNA*^RUS^* transcriptome profiles at both time points (Fig EV3C).

Next, we determined differentially expressed genes (Fig EV3D, Table EV2) and analyzed enriched gene ontology (GO) classifications (Mi *et al*, 2013) among the up- and down- regulated genes, separately for the two time points. Depletion of *RUS* massively affected the transcriptome: on day 5, 4978 genes (24%) were transcribed at elevated levels under reduced *RUS* levels and 4586 genes (22%) were repressed (Fig EV3D). The expression changes were even more profound on day 7, when 6623 genes (30%) and 6456 genes (29%) were up- or downregulated, respectively.

In agreement with the observed increase of apoptosis upon *RUS* depletion, we found the GO annotations associated with ’cell death’ and ‘apoptosis’ (represented by ‘positive regulation of apoptosis’ in Fig 3A) enriched among the induced genes on both days 5 and 7, exemplified by genes encoding, Bak1, and Foxo3. Figure 3B shows these genes among the 50 most deregulated genes enriching for the GO annotations: ’cell-death’, ’neurogenesis’, ’cell-cycle’ and ’microtubule-based process’. Annotations represented by GO classifications ‘cell cycle and ‘microtubule-based process’ (Fig 3A) were most significantly enriched among the downregulated genes on both days, in support of the reduced BrdU incorporation (Fig EV2E) and indicative of proliferation arrest (Fig 3A,B). Interestingly, genes with GO annotations relating to ’neurogenesis’ and ’neuron differentiation’ were mildly enriched among the downregulated on day 5, but strongly enriched among the induced genes on day 7 (Fig 3A,B). Of note, at this level of analysis direct and indirect effects cannot be distinguished.

**Figure 3.**
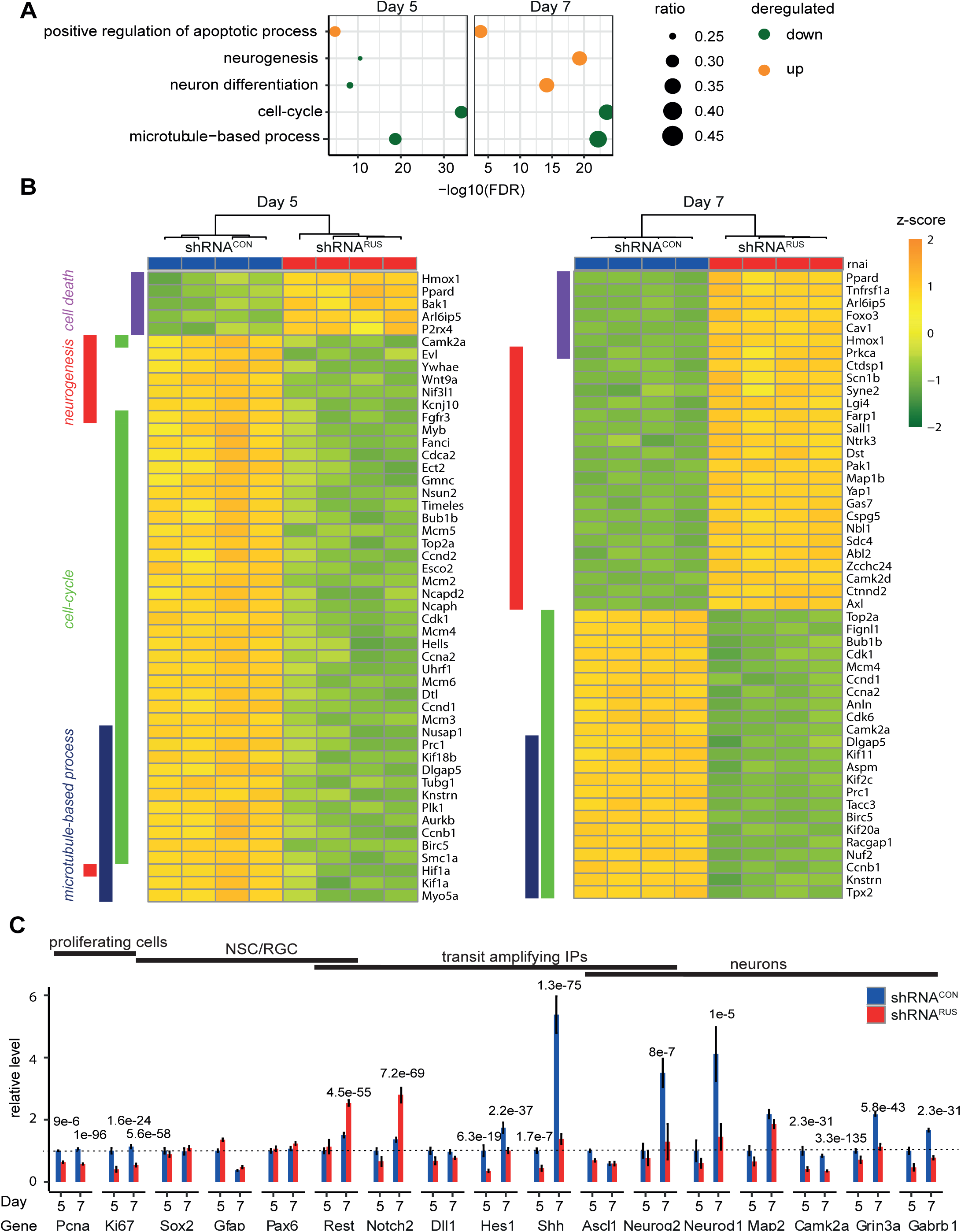
Transcriptome changes upon depletion of *RUS*. **A.** Enriched gene ontology (GO) classifications among genes down-regulated (blue) or up-regulated (orange) upon *RUS* depletion at days 5 and day 7 of culture, as indicated. Circle size indicates the number of deregulated genes compared to the total number of genes enriched in the respective GO annotation (100% =1). **B.** Heatmap showing the top 50 deregulated genes enriching for the GO annotations ’cell-death’, ’neurogenesis’, ’cell-cycle’ and ’microtubule-based process’ on day five (left) and day 7 (right) of culture. Note that these are different genes. The genes were sorted by GO annotations and difference between shRNA*^CON^*and shRNA*^RUS^*,. **C.** Expression levels of the indicated marker genes on day 5 and day 7 of culture in control (shRNA*^CON^*, blue) and knockdown (shRNA*^RUS^*, red) cells were determined by RNA-seq (TPM values were normalized to those of the control cells on day 5. Error bars show the standard deviation)

To explore the effects of *RUS* depletion in our RNA-seq data in more detail, we determined the read counts of several prominent genes that characterize the *in vitro* differentiation process (Fig 3C). We assessed the proliferation state (*Pcna* and *Ki67*), the NSC/RGC markers *Sox2*, *Pax6* and *Gfap* as well as the neuronal markers *Neurog2*, *Neurod1*, *Map2, Camk2a*, *Grin3a*, and *Gabrb1*. In addition, we focused on the Notch1/2 and sonic hedgehog (Shh) signaling pathways regulating the expansion of RGCs and transit-amplifying intermediate progenitor cell populations. *Notch1/2,* its ligand *Dll1* and their downstream effectors *Hes1, Neurog2, and Ascl1* form an oscillatory network that regulates RGC cell renewal (Hatakeyama & Kageyama, 2006; Wang *et al*, 2016; Ivanov, 2019; Sueda & Kageyama, 2019). We also included *Rest* as a transcriptional repressor of neuro-specific genes which helps to maintain the neural stem cell state (Schoenherr & Anderson, 1995; Mukherjee *et al*, 2016).

Our RNA-seq analysis confirmed that the proliferative markers *Pcna* and *Ki67* were robustly downregulated on both day 5 and day 7 (Fig 3C). The NSC/RGC markers *Sox2, Pax6, Gfap* were less affected. However, the substantially reduced expression of the neuronal cell fate commitment markers *Hes1*, and *Shh* as well as of the neuronal markers: *Neurog2*, *Neurod1*, *Camk2a*, *Grin3a*, and *Gabrb1* confirmed our earlier notion that depletion of *RUS* compromises neuronal differentiation. Of note, the expression of those genes that are most strongly induced during neurogenesis between days 5-7 (i.e., *Shh*, *Neurog2*, and *Neurod*) was most strongly affected by *RUS* depletion (Fig 3C). The increased expression of Notch2 is consistent with the observed maintenance of NSC/RGC markers, the reduced expression of cell cycle genes as well as genes involved in neurogenesis (Engler *et al*, 2018; Mase *et al*, 2021). The induction of *Rest* at day 7 suggests a mechanism involving chromatin regulation.

We conclude that *RUS* is required for efficient proliferation and for differentiation of neuronal precursor cells in this *in vitro* system. The concomitant inhibition of cell proliferation (and hence cell renewal) and neurogenic differentiation may leave neuronal progenitor cells with conflicting signals that trigger apoptosis. The observation that at day 7 the most deregulated genes with annotated GO term ‘neurogenesis’ are activated upon *RUS* depletion (Fig 3B) prompts the speculation that *RUS* may be involved in the repression of transcription. Again, direct and indirect effects cannot be distinguished at this point.

### RUS associates with chromatin of key neurodevelopmental genes

As a first step towards defining the mechanism through which *RUS* regulates gene expression, we determined the subcellular localization of *RUS*. After two days in culture, cells were fractionated into the cytoplasm, nucleoplasm and chromatin. RT-qPCR analyses showed that *RUS* is enriched in the chromatin fraction, similar to the splicing-associated lncRNA *MALAT* (Fig 4A). An RNA-FISH (fluorescence-*in-situ*-hybridization) experiment confirmed the nuclear localization (data not shown).

**Figure 4:**
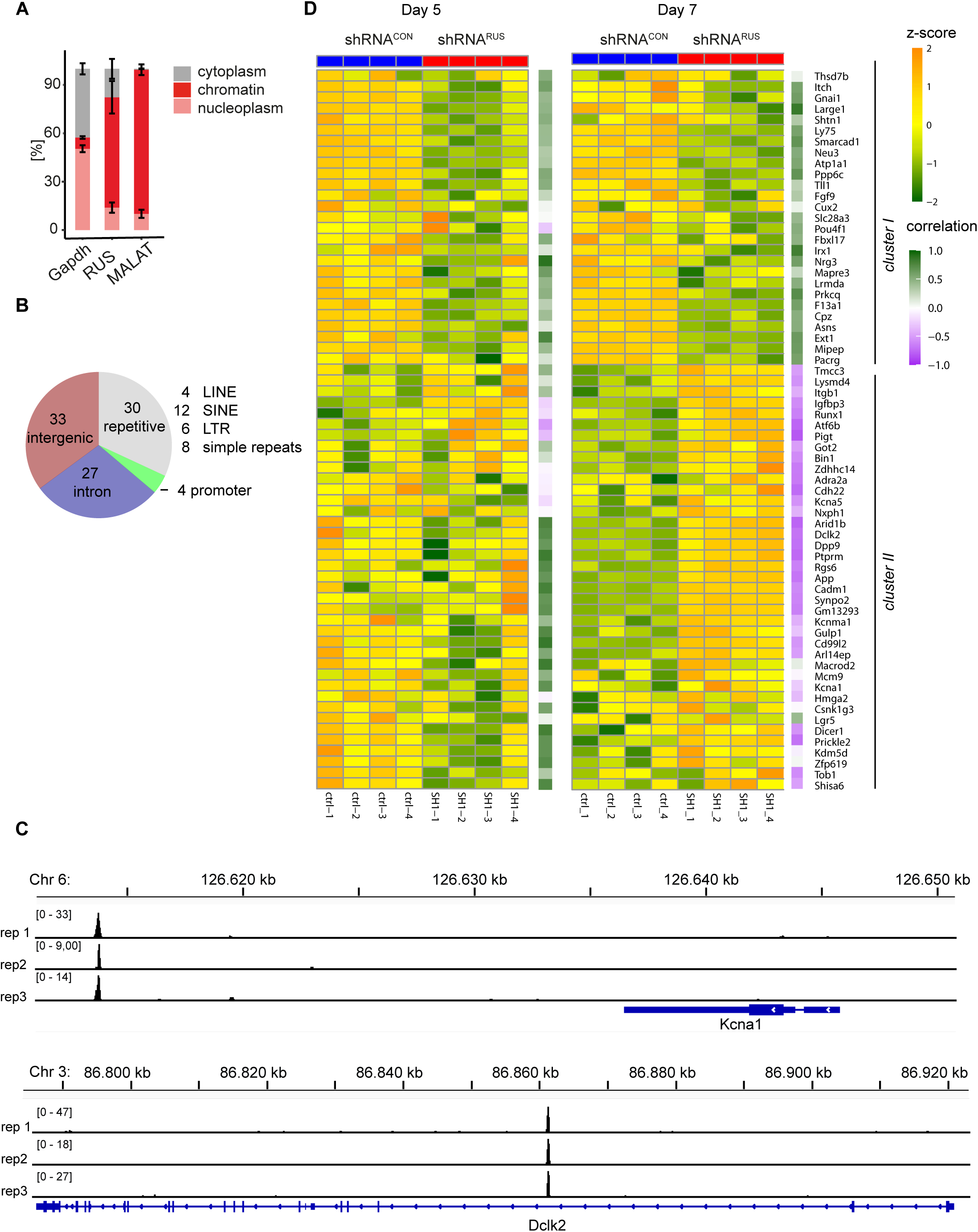
Localization of *RUS* to chromosomal sites. **A.** Subcellular localization of *RUS*. NSCs were differentiated for two days and then fractionated into the sub-cellular compartments cytoplasm, nucleoplasm and chromatin. *RUS* was detected by RT-qPCR along with *Gapdh* mRNA (cytoplasmic marker) and *MALAT* (nuclear marker). Error bars: standard deviation of 3 independent experiments. **B.** Genome annotation of the 94 high-confidence *RUS* ChIRP locations. SINE: short interspersed nuclear element; LINE: long interspersed nuclear element; LTR: long terminal repeat. **C.** Browser view of two examples of *RUS* localization close to relevant neurogenic genes. The *RUS* ChIRP tag density of the three replicates is plotted I separate tracks in the genomic regions of the *Kcna1* (top) and *Dclk2* (bottom) genes. For orientation, the respective chromosomal regions are displayed above and the gene models below the traces. **D.** Heatmap showing the expression changes of 66 *RUS* putative target genes upon *RUS* depletion (shRNA*^RUS^*, red) or in control cells (shRNA*^CON^*, blue) on days 5 and 7. Replicate identifiers are indicated below the columns. Genes were hierarchically clustered using Euclidean distance based on their combined expression on both days. This yields two clusters depending on whether genes are activated or repressed upon *RUS* depletion. The gene names are indicated to the right of the 7- day heatmap. The purple-green code to the right of each individual heatmap indicates the degree of correlation between *RUS* and putative target gene expression.

To explore whether *RUS* localizes to specific chromosomal regions like other regulatory lncRNAs, we applied the ChIRP (<u>Ch</u>romatin Isolation by <u>R</u>NA <u>P</u>urification) methodology (Chu *et al*, 2011). Cells were harvested at day 7 of differentiation and *RUS* was isolated by hybridization with two independent probe sets (‘odd’ and ‘even’). The experiment was done in biological triplicate. All three isolations effectively retrieved *RUS* (approximately 30% of input) and strongly enriched *RUS* over control RNAs *TBP mRNA*, *MALAT* and *XIST* (Fig EV4A). Between 157 to 203 peaks were scored in individual experiments, of which 129 (67%, Fig EV4B) overlapped in all three experiments (Table EV3).

Although we considered only peaks enriched by both probe sets, several enriched genomic sites contained sequences with similarity to one of the used oligonucleotide probe sequences. After removing them, 94 high-confidence putative *RUS* binding sites remained for further analysis (for simplicity called ‘*RUS* binding sites’ below). Genomic annotation revealed that 4 of them (4.3%) mapped to promoters, but the majority predominantly localized to intergenic (35.1%) or intronic (28.7%) regions, compatible with long-range regulatory elements. About a third of the locations mapped close to degenerate repetitive elements of various types, such as LINEs (4.2%), SINEs (12.8%), LTR (6.4%) and simple repeats (8.5%) (Fig 4B). Gene ontology analysis of the active genes next to *RUS* binding sites yielded an enrichment of the terms ‘forebrain development’, ‘neurogenesis’ and ‘generation of neurons’. Among those are the genes encoding the microtubule stabilizing protein Dclk2 and the potassium voltage-gated channel Kcna1 (Fig 4C, two further tracks: *Arid1b* and *Bin1* in Fig EV4C). Both genes play a pivotal role in neuron differentiation (Shin *et al*, 2013; Chou *et al*, 2021).

Following the hypothesis that *RUS* binding to chromatin is involved in regulating near-by genes, we determined the expression changes of genes residing next to *RUS* binding sites (referred to as ‘putative target genes’ henceforth) using the RNA-seq data of *RUS* knockdown samples. Of the 94 putative target genes, 66 were robustly expressed in differentiating NSC (Fig 4D). The number of genes that changed their expression increased from day 5 to day 7 (54% and 77% of genes with altered expression, respectively), in line with the increase of *RUS* expression between days 5 and 7 of differentiation (Table EV3).

Hierarchical clustering of expression separates putative target genes into two distinct clusters (Fig 4D). Cluster I contains genes significantly downregulated on both days, while cluster II represents genes with enhanced expression, predominantly on day 7. The heat map shows several cluster II genes with reduced expression on day 5 after *RUS* depletion. Since *RUS* depletion was less effective on day 5, we calculated the overall correlation of *RUS* expression and its putative target genes (Fig 4D, purple-to-green boxes to the right of heat maps). If we assume direct effects of *RUS* binding on target gene expression, we expect a positive correlation of genes with reduced expression with *RUS* depletion (essentially genes in cluster I) and a negative correlation of genes with enhanced expression upon *RUS* depletion (predominantly cluster II genes on day 7). This is indeed largely the case (Fig 4D). Remarkably, the expression of genes that are repressed on day 5 and activated on day 7, for example *Arid1b*, *App*, and *Kcna1* (Fig EV4D), correlates positively on day 5 and negatively on day 7 with *RUS* expression, in support of a direct effect of RUS on close-by genes. Our results thus suggest that RUS may mediate both, activating and repressive regulation.

### RUS interactors suggest epigenetic regulatory mechanisms

LncRNAs usually elicit their gene regulatory effects through interacting effector proteins. To explore how *RUS* may mediate both, activating and repressive functions, we sought to identify *RUS*-binding proteins. When mouse and human *RUS* sequences are compared, a remarkable degree of conservation of exon 1 stands out (Fig 1C). Because such conservation may be indicative of important functional interactions, we compared interactors of complete *RUS* with a 5’-deleted RNA *(Δ5’-RUS*), lacking exon 1. Both RNAs were tagged with 5 MS2 stem-loop structures at the 3’ end, enabling affinity purification via binding to MS2-binding protein (MS2BP) (Johansson *et al*, 1997; Zhou *et al*, 2002; Tsai *et al*, 2011).

Because differentiating NSCs cannot be obtained in sufficient amounts for RNA-affinity purification, we established an RNA-affinity purification protocol using the well-established Neuro2A cell line*. RUS* is normally not expressed in these cells and so our experiment identifies potential protein interactors that are not relevant in these cells. To assure an equivalent expression of both RNAs, we first generated a Neuro2A derivatives by inserting an FRT recombinase site into the genome through lentiviral transduction. These clonal cells were then transfected with FRT-flanked *RUS* expression constructs along with a flipase expression plasmid (Andrews *et al*, 1985; Sauer, 1994; See *et al*, 2002)). Clones containing integrated *RUS* expression cassettes were expanded and analyzed. These clones express comparable levels of either full-length *RUS* or *Δ5’-RUS*.

Lysates of *RUS*- and Δ5’-*RUS*-expressing cells were incubated with recombinant MS2- binding protein (MS2BP), which in turn was tagged with a maltose-binding protein (MBP) (see scheme in Fig 5A). MS2BP-bound RNA was retrieved by absorption of MBP to amylose beads, captured proteins were eluted with RNAse A treatment and identified by LC-MS, using label-free quantification (LFQ) (Cox & Mann, 2009).

**Figure 5:**
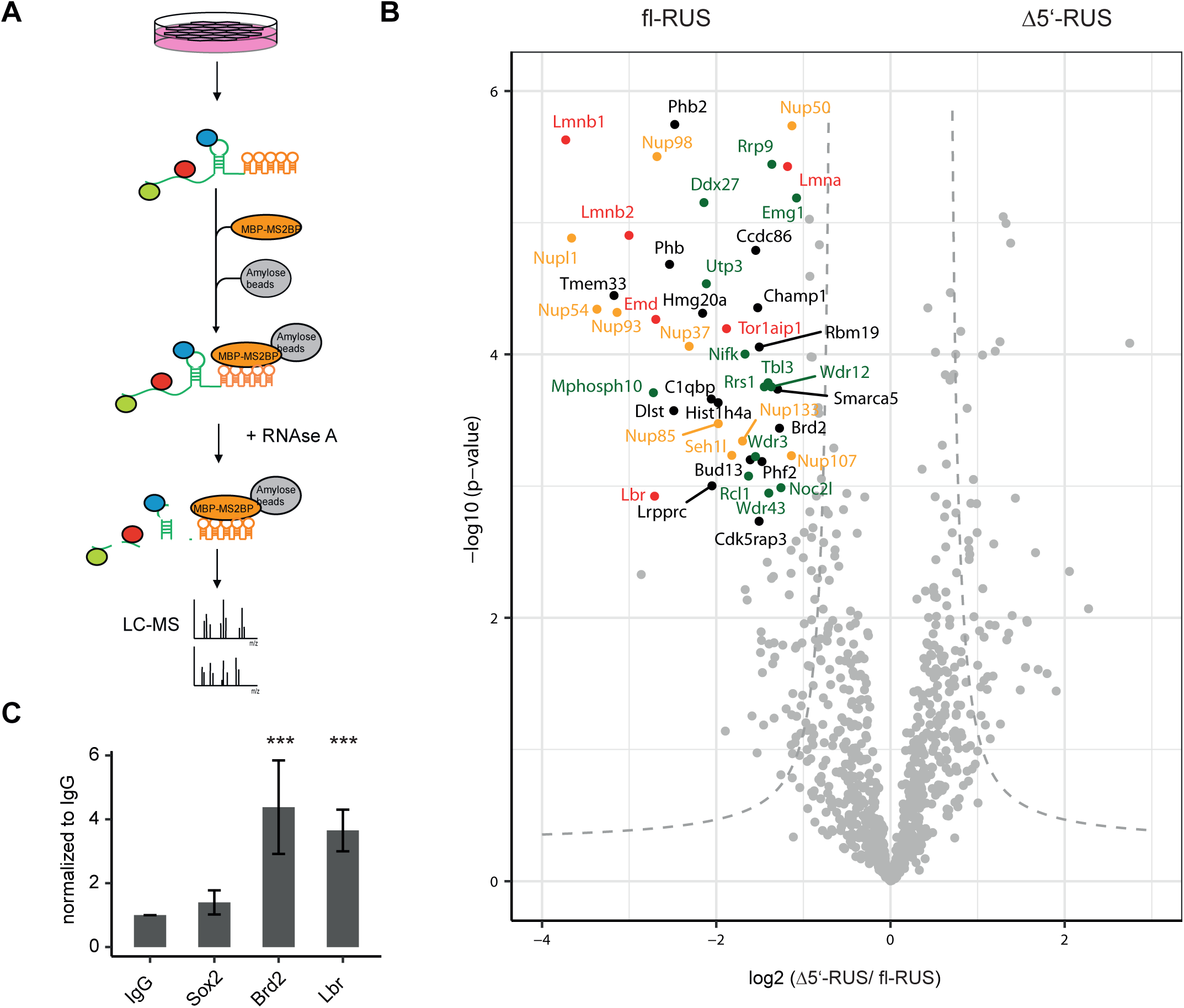
RUS interacts with components of the nuclear pore, -lamina, and nucleolus. A. Schematic overview of the affinity purification of *RUS*-interacting proteins (colored spheres). *RUS* RNA (green), tagged with 5 MS2 stem-loop structures (orange) is stably expressed in Neuro2A cells. The RNA is affinity-purified by binding to MS2BP- maltose binding protein on an amylose resin. For details, see text. B. Volcano plot showing affinity-purified nuclear proteins that bind differentially to full-length *RUS* (left) or a *RUS* RNA from which exon 1 was deleted *(Δ5’-RUS*). Proteins with a change greater than 2 and a p-value smaller than 0.002 are considered robust interactors and annotated by their gene name. The dashed gray hyperbolic curves depict a permutation-based false discovery rate estimation (P = 0.05; s0 = 1). Some proteins are color-coded: proteins of the nuclear lamina (red), nuclear porins (orange) and nucleolar proteins (green). C. Quantitative RT-PCR analysis of *RUS* co-immunoprecipitated with antibodies against Sox2, Brd2, Lbr and control IgG from differentiating NSCs. Error bars show the standard deviation (*** p < 0.01 compared to IgG purification).

Full-length *RUS* enriched many more proteins in comparison to Δ5’-*RUS* (Fig 5B, Table EV4). While we cannot exclude that this is due to the increased size of the *RUS* RNA, this seems unlikely given the size difference of 912 (*RUS*) versus 679 nucleotides (*Δ5’-RUS).* Proteins with a fold-change greater than 2 and a p-value smaller than 0.002 were considered robust and specific binders. Only 9 proteins were purified selectively along with Δ5’-*RUS.* By contrast, 49 proteins were enriched by co-purification with the full-length construct and therefore considered exon 1-specific interactors (Table 1, EV4). Among them, Phb, Phb2, Tor1aip1 and Utp3 were purified exclusively by the full-length *RUS* RNA.

**Table 1:**
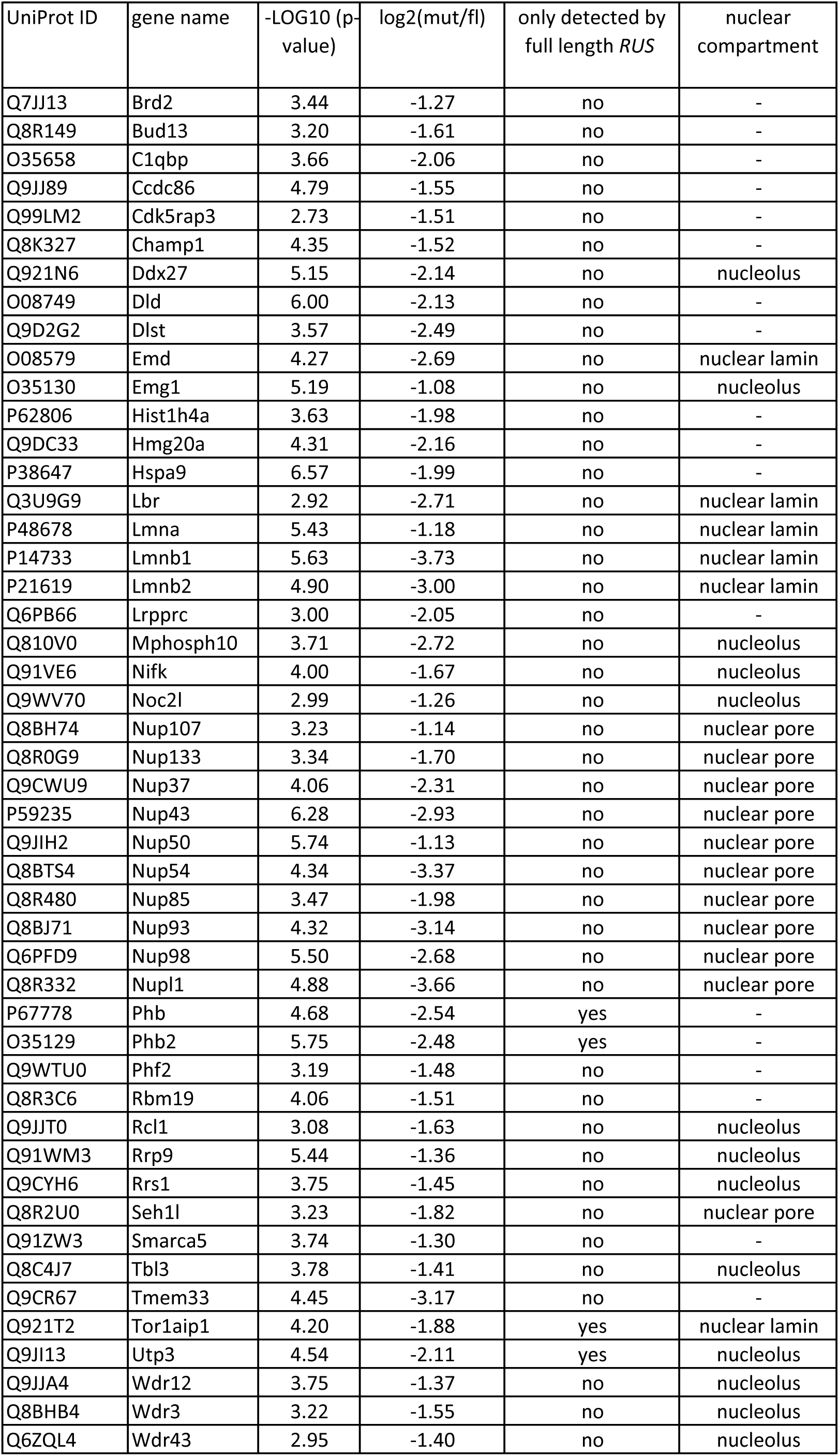
Table includes affinity-purified nuclear proteins that bind more than full length *RUS* (p-value < 0.002, log2(mut/fl RUS) < -1) and the localization to nuclear compartments as nucleolus, nuclear lamin and nuclear pore. Table highlights whether a protein was identified by full-length *RUS* only.

**Table 2:**
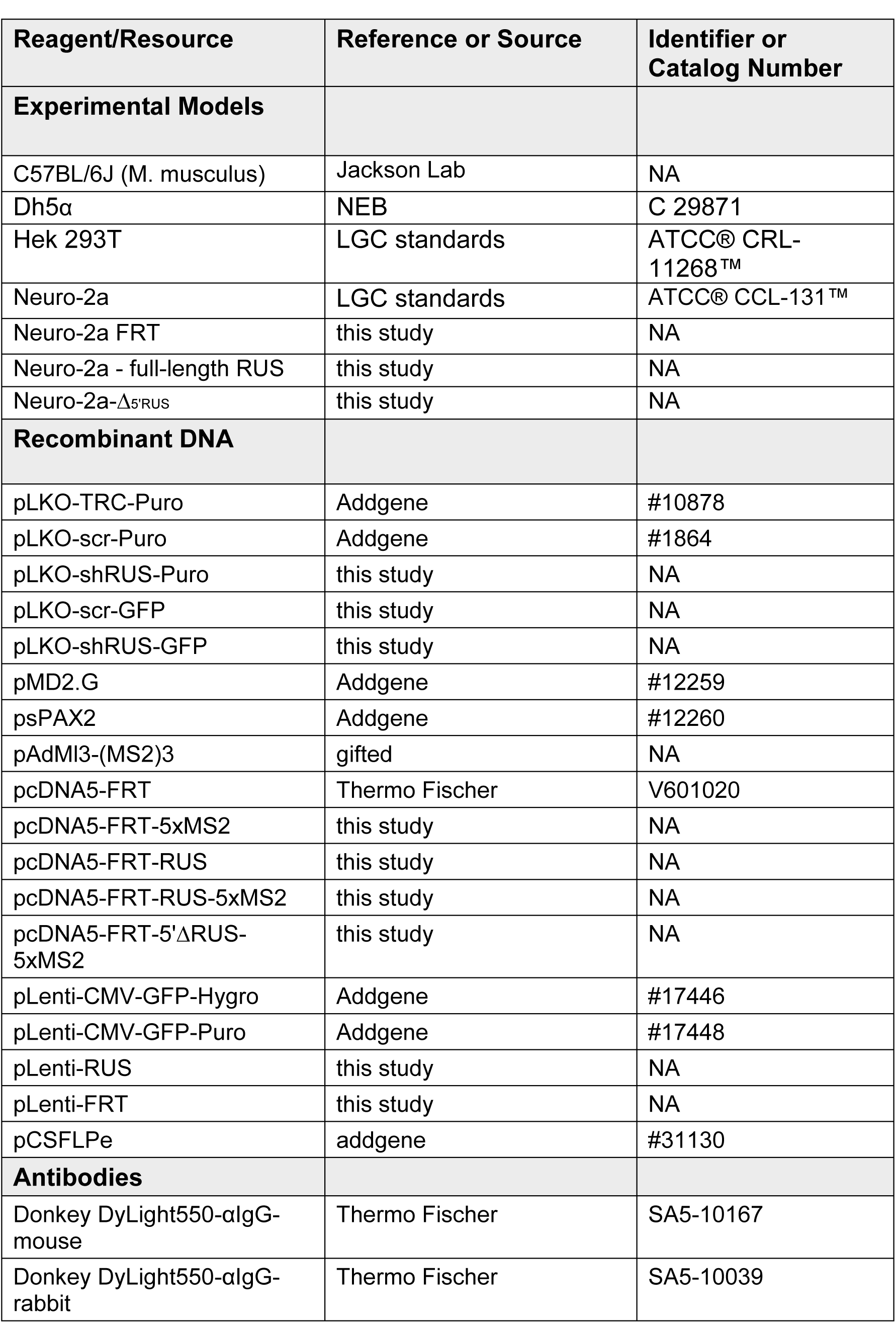
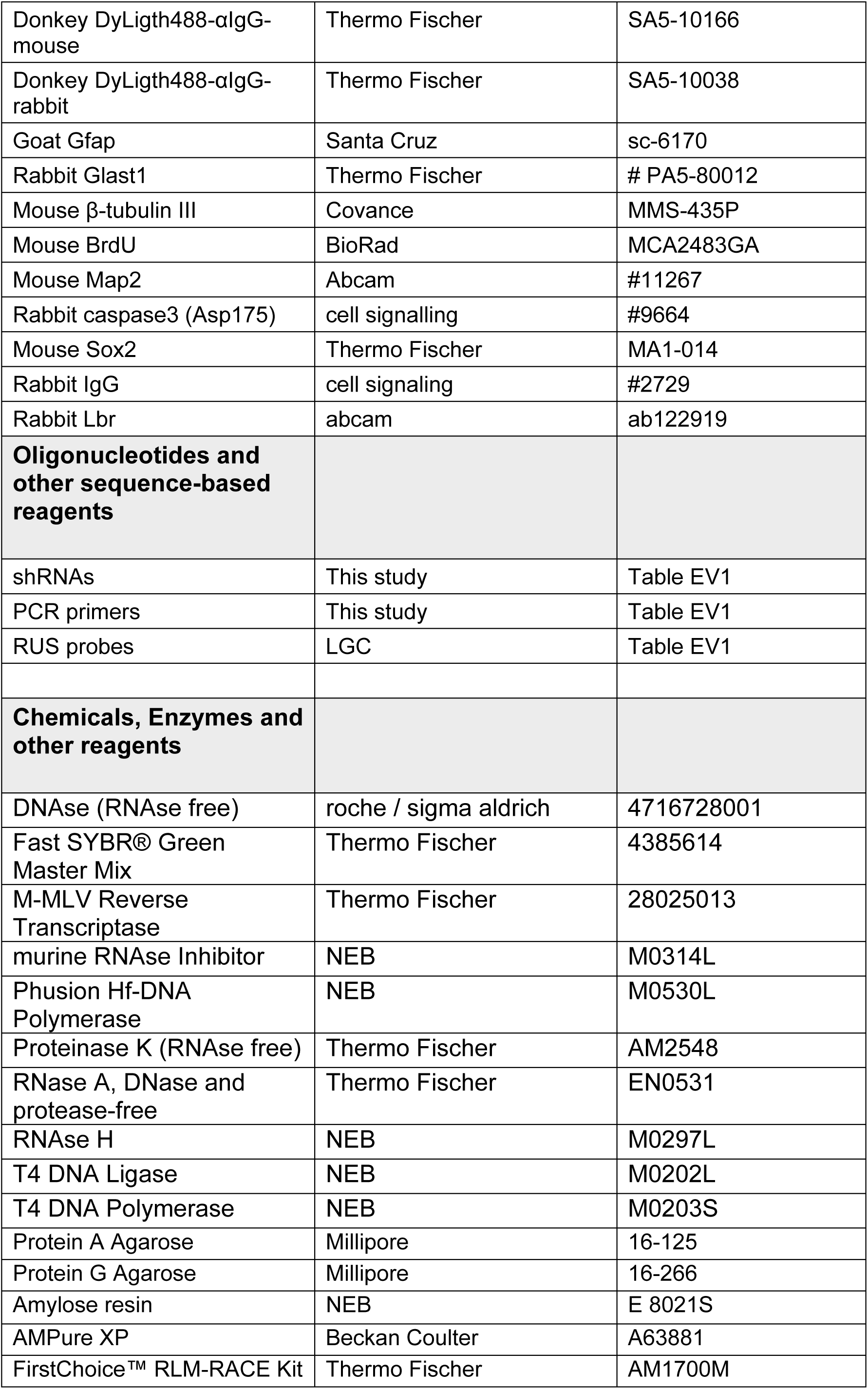
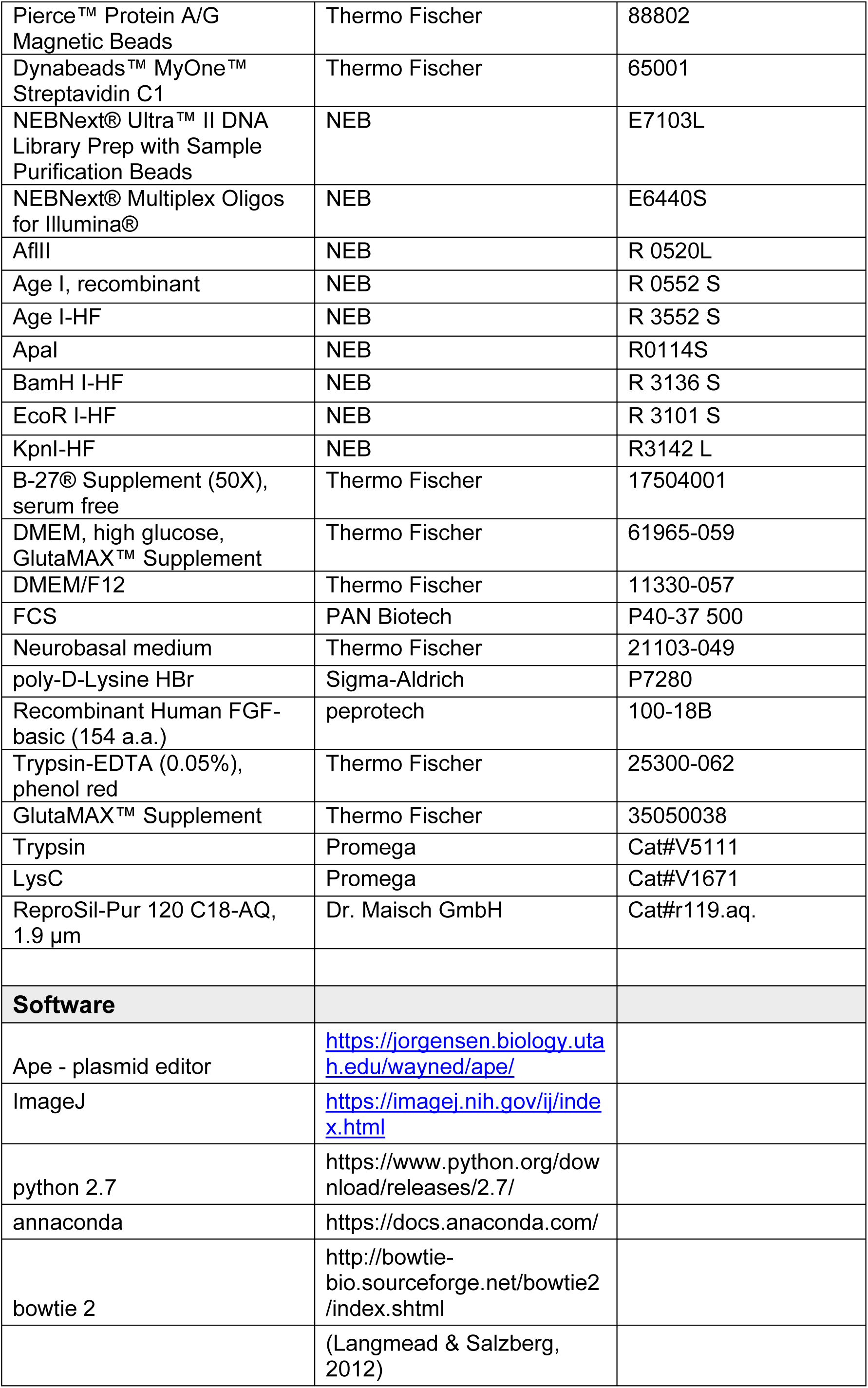
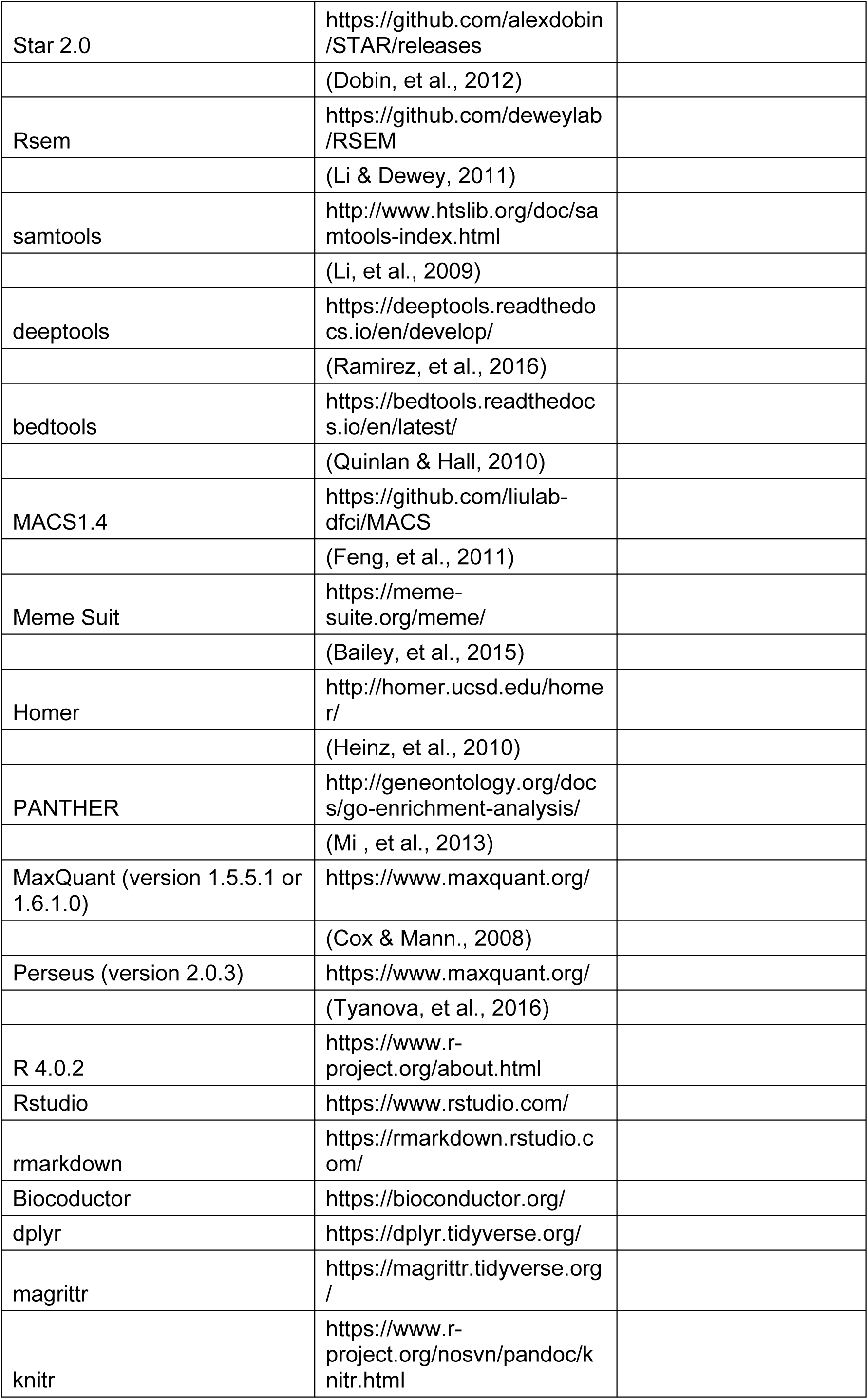
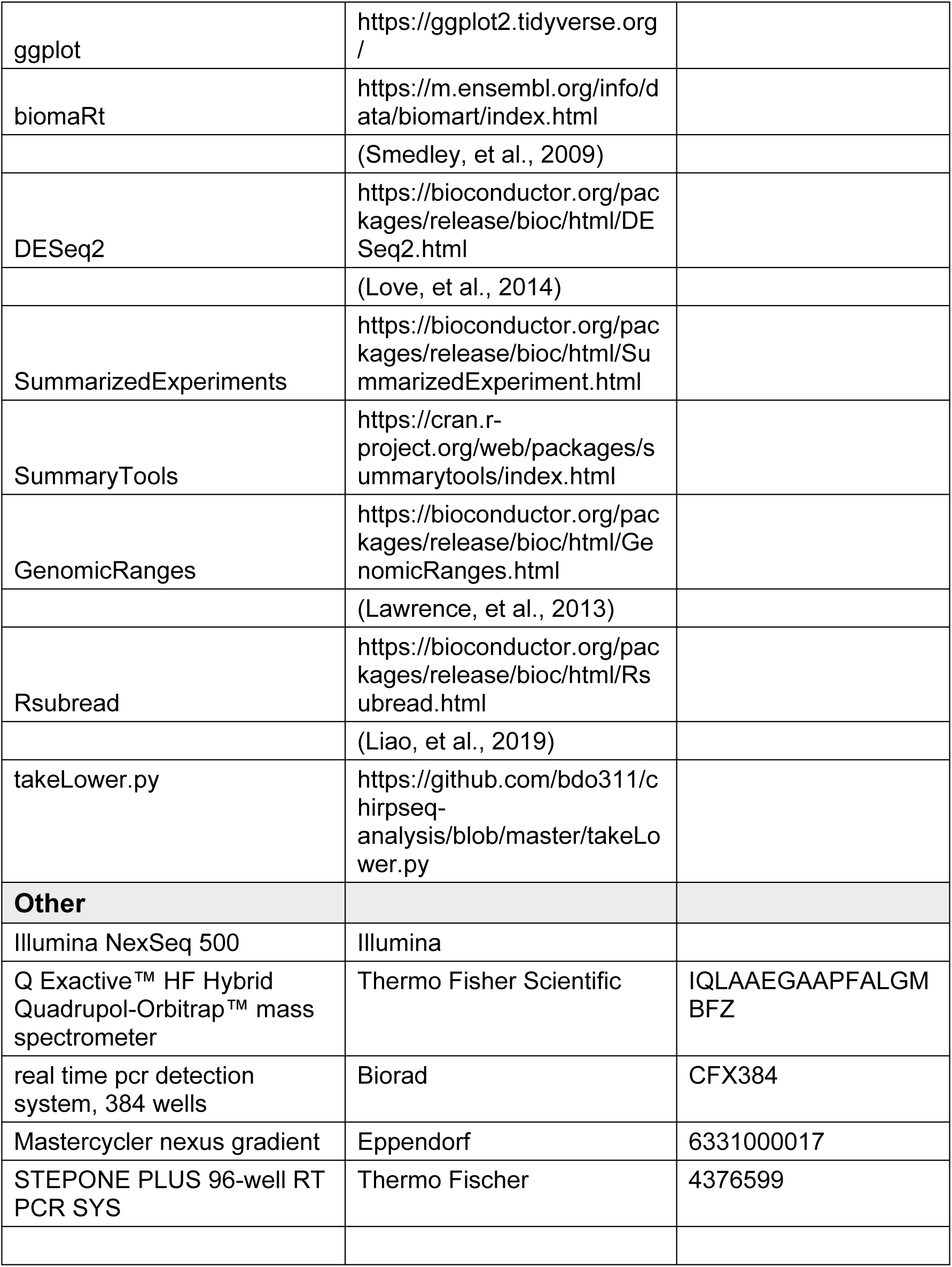
Reagents and Tools table

Phb and Phb2 correspond to the prohibitin complex, a mitochondrial regulator with neuroprotective functions and nuclear co-repressor of cell cycle-regulated genes (Koushyar *et al*, 2015).

Further, we find numerous components of the nuclear periphery, most prominently subunits of the nuclear pore complex (Nupl1, Nup37, Nup43, Nup50, Nup54, Nup85 Nup93, Nup98, Nup107, Nup133, and Seh1l orange in Fig 5B) and several constituents of the nuclear lamina: emerin (Emd), lamins A, B1 and B2 (Lmna, Lmnb1, Lmnb2), lamin B receptor (Lbr) as well as the lamin A/B binding protein Tor1aip1 (red in Fig 5B).

Further, *RUS* exon 1 retrieved many nucleolar proteins (Ddx27, Emg1, Mphosph10, Noc2l, Nifk, Rcl1, Rrp9, Rrs1, Tbl3, Utp3, Wdr3, Wdr12 and Wdr43 green in Fig 5B) and some interesting chromatin regulators (e.g. the bromodomain protein Brd2, the chromatin constituent Hmg20a, the nucleosome remodeling ATPase Smarca5, the lysine demethylase subunit Phf2 and the RNA helicase Ddx54).

The finding of robust interaction of *RUS* with nuclear pores and the lamina suggest well-established epigenetic regulatory mechanisms (to be discussed below). Binding of lncRNA *Xist* to Lbr has been suggested to tether the inactive X chromosome to the nuclear envelope, which forms a silent compartment (Chun-Kan *et al*, 2016). To validate the binding between *RUS* and Lbr we returned to our NSC differentiation model. Nuclear extracts were prepared from cells harvested at day 7 of differentiation. Lbr was immunoprecipitated and co-precipitated RNA quantified by RT-qPCR. *RUS* was retrieved 3.7-fold more by comparison to an anti-IgG purification (Fig 5C). Parallel reactions confirmed the selective interaction of Brd2 with *RUS*, while Sox2 served as a control.

In summary, our data support the idea of the long, noncoding RNA *RUS* as a crucial regulator of the neurogenic gene expression program through epigenetic mechanisms.

## Discussion

### The lncRNA RUS is required to execute the neurogenic program

Our study presents a first functional characterization of the lncRNA *LINC01322,* which we term *RUS* (for <u>R</u>NA <u>u</u>pstream of *<u>S</u>litrk3*). Like other neurogenic lncRNAs, *RUS* is well conserved in mammals by sequence and synteny next to the neurodevelopmental gene *Slitrk3*. It is predominantly expressed in neural tissues. Although the RNA bears some coding potential, we did not detect any of the theoretically encoded peptides. *RUS* associates with chromatin at specific sites in the vicinity of neurodevelopmental genes and interacts with several proteins involved in epigenetic gene regulation, suggesting that *RUS* acts as lncRNA.

Transcriptome analyses revealed that sh-mediated depletion of *RUS* results in massive gene expression changes. In fact, approximately half of all genes were affected to a certain degree. The responses were equally divided between gene activation and repression and were modulated during the 7 days of differentiation. This finding is interesting, since most lncRNA studied so far either mediate activation or repression (Rinn & Chang, 2020; Statello *et al*, 2021). Although indirect effects cannot be excluded yet, the fact that we found epigenetic activators and repressors bound to *RUS* exon 1 in pulldown experiments, supports the idea that *RUS* may mediate gene activation and repression in a highly context-dependent manner. Conceivably, *RUS* may function through diverse mechanisms, as emerges for the HOTAIR RNA (Price *et al*, 2021).

On day 5 of differentiation, reduced *RUS* levels correlate with reduced expression of many genes involved in neurogenesis and cell cycle, suggesting that the lncRNA promotes target gene expression to enable amplification of intermediate precursor cells and NSC differentiation. This is in line with the observation that *RUS* is expressed in hESC-derived LRGs (Ziller *et al*, 2015).

*RUS* is most highly expressed in the adult hippocampus, in which neurogenesis still occurs (Eriksson *et al*, 1998). Adult neurogenesis relies on expanding transit-amplifying IPs maintained by *Shh* expression (Antonelli *et al*, 2018) and differentiation by increased *Neurog2* expression (Galichet *et al*, 2008). At day 7 of our differentiation time course, *Shh*, *Neurog2*, and *NeuroD1* are among the most repressed genes upon *RUS* depletion. In addition, we found a reduced expression of several subunits of glutamate and GABA receptors, such as *Grin3a* and *Gabrb1,* which are predominantly expressed in neurons.

We propose that *RUS* depletion locks neuronal precursors in an intermediate state towards neuronal differentiation, with arrested cell cycle. The activation of pro-apoptotic genes may result from perturbed cell identity.

### Potential mechanisms of RUS-mediated gene regulation

Given the diverse and presumably very site-specific effects of *RUS* function, we can only speculate about potential mechanisms. Our stringent ChIRP approach revealed a very consistent set of *RUS* interactions with a limited number of high-confidence chromatin loci. The localisation of binding sites predominantly in introns and intergenic regions argue for long-range regulation. Considering that the RNA is not highly expressed, we speculate that its range of activity may be limited to the genes in the vicinity of tethering sites (Engreitz *et al*, 2016).

Remarkably, most of the genes closest to a *RUS* binding site were expressed in differentiating NSCs and changed their expression state upon *RUS* depletion. For example, *RUS* binds in the genome next to genes essential for cell cycle and neuronal differentiation, such as *Fgf9, Mapre3, and Ppp6c, Arid1b, Dclk2, and Kcna1.* The expression of these critical genes is affected by *RUS* depletion. Furthermore, *RUS* binding sites can be observed in introns of the E3 ubiquitin ligase genes *Itch* and *Fbxl17*. Itch ubiquitinates Notch proteins for degradation to turn off Notch signaling (Chen *et al*, 2021). *Fbxl17* plays a pivotal role in Shh signaling by degrading Sufu to enable the translocation of Sufu-sequestered transcription factors to the nucleus (Raducu *et al*, 2016). Consequently, reduction of both factors after *RUS* depletion resulted in increased Notch signaling and reduced Shh signaling, consistent with our RNA-Seq data. Notch signaling is important for maintaining the active or quiescent neural stem cell state by preventing neuronal differentiation (Sueda & Kageyama, 2019). Shh signaling regulates proliferation of neural precursors (Yao *et al*, 2016). By activating both genes *RUS* facilitates proliferation and ensures proper differentiation of neural precursor cells.

LncRNA often work by recruiting epigenetic regulators to locally concentrate them at target chromatin (Markaki *et al*, 2021). Our RNA-affinity purification relies on protein-*RUS* interactions formed under physiological conditions in intact cells and purifying complexes under native conditions. Because we wished to identify proteins interacting with the conserved exon 1 of *RUS*, we monitored the differential binding to RNA containing or lacking this sequence. This is a stringent approach, because functionally meaningful proteins may well (and are indeed likely to) bind to the remainder of *RUS* as well, but they are not discussed here (but see Table EV4). In the following, we discuss hypothetical scenarios, in which *RUS* recruits regulatory functions to chromosomal target loci. It is also possible that *RUS* sequesters the factors in competition with other interactors, which would have opposite effects on gene regulation compared to recruitment scenarios (Xi *et al*, 2022).

Among the proteins purified by full length *RUS* only, the prohibitin complex (consisting of Phb and Phb2) stands out. Prohibitin has functions in several cellular compartments, including mitochondria and nuclei (Wang *et al*, 2002; Fusaro *et al*, 2003; Rajalingam & Rudel, 2005; Koushyar *et al*, 2015). Prohibitin has been termed an oncogene, as it promotes proliferation and dedifferentiation in neuroblast cells (MacArthur *et al*, 2019) and a tumour suppressor gene, since it was shown to inhibit the cell cycle by repressing E2F-regulated genes via recruitment of the retinoblastoma protein and histone deacetylases (Wang *et al*, 2002). It is tempting to speculate that tethering the Phb complex to chromatin contributes to inhibition of proliferation and activation of apoptosis.

Strikingly, the RNA pulldown retrieved numerous proteins of the nuclear envelope. We scored 6 constituents of the nuclear lamina, including three types of lamins and lamin B receptor (Lbr). The inner nuclear membrane assembles a well-known repressive compartment to which inactive heterochromatin is tethered. These lamina-associated domains may be constitutive or facultative (van Steensel & Belmont, 2017). Conceivably, *RUS* mediates tethering of genes destined to be silenced to the lamina, where they acquire heterochromatic features. Such a scenario has precedent in the finding that the lncRNA *XIST* promotes X chromosome inactivation in female cells by tethering the target chromosome to the nuclear envelope via Lbr (Chun-Kan *et al*, 2016).

Repressive heterochromatin is also found at the surface of nucleoli (Kind *et al*, 2013; Vertii *et al*, 2019). Remarkably, we found 13 nucleolar proteins enriched specifically by *RUS* exon 1, which further supports the speculation that *RUS* partitions genes into silencing compartments. However, some of the retrieved nucleolar proteins also have nuclear functions. For example, NOC2L (NOC2 Like Nucleolar Associated Transcriptional Repressor, a.k.a. NIR) associates with p53 in the nucleus to repress a subset of p53-target genes, including p21, by inhibition of histone acetylation (Hublitz *et al*, 2005). Interestingly, the exon 1 interactor NIFK (also a nucleolar protein with nuclear functions) also cooperates with p53 to silence the p21 promoter during checkpoint control (Takagi *et al*, 2001). Apparently, *RUS* also contributes to p21 silencing since the gene gained activity upon depletion of the lncRNA. Similarly, the exon-1 interactor Cdk5rap3 activates p53 activity by repressing its degradation by Hdm2 (Wang *et al*, 2006). Such a scenario provides a plausible and testable hypothesis for the observed cell cycle arrest at reduced *RUS* levels.

In addition to constituents of the nuclear lamina, we found 11 nuclear pore components (Nup11, Nup37, Nup43, Nup50, Nup54, Nup85, Nup93, Nup98, Nup107, Nup133 and Seh1l) among the exon 1 interactors. In addition to nuclear transport, the nuclear pore complex plays an important role in transcriptional regulation and cell identity, apparently by generating a microenvironment that fosters epigenetic regulation of associated genes (Pascual-Garcia & Capelson, 2021). In *Drosophila*, Nup93 is associated with genes repressed by the polycomb complex and is required for efficient repression (Gozalo *et al*, 2020).

By contrast, three nucleoporins bound *RUS* are predominantly associated with transcriptional activation. Nup98 acts as anchor point for enhancer (Pascual-Garcia *et al*, 2017) and activates transcription by recruiting the Wdr82-Set1A/COMPASS complex to regulate H3K4 trimethylation (Franks *et al*, 2017). Similarly, Nup107 and Seh1l activate transcription by assembling transcription factor (TF) complexes at the nuclear pore (Liu *et al*, 2019). It is tempting to speculate that *RUS* may mediate facultative association of gene loci with the nuclear periphery, which would then be subject to regulation of the corresponding microenvironment. This may initially involve an initial transcriptional activation to execute the differentiation programme. The subsequent compartmentalization of chromosomal loci into a repressive environment may serve to terminally silence cell cycle genes in mature neurons.

The exon 1 interactor HMG20A (a.k.a. iBraf) is known to antagonize repressive LSD1-REST complexes. Since LSD1-REST-dependent H3K4 demethylation represses neuronal genes, HMG20A action promotes neuronal differentiation (Ceballos-Chávez *et al*, 2012; Garay *et al*, 2016). The interaction of *RUS* with HMG20A, therefore, likely affects neuronal differentiation, but whether the outcome is positive (through recruitment) or negative (through squelching) remains to be explored. Of note, REST expression increases upon *RUS* depletion, consistent with the observed inhibition of neurogenesis.

In summary, our mapping of putative target genes and *RUS* interactors are compatible with a range of testable, hypothetical and not mutually exclusive scenarios that may explain the observed change in phenotype and gene expression upon *RUS* depletion during differentiation of NSCs. We propose that *RUS* may be involved in several aspects of the neurogenic program in a highly context-dependent manner, including amplification of precursor cells and terminal neuronal differentiation.

## Materials and Methods

### Cultivation and differentiation of primary neural stem cells

The isolation of cortical embryonic stem cells from E15-E16 murine cortices was approved by the animal welfare committees of LMU and the Bavarian state. Cortices were dissected from pooled mixed-sex embryonic brains, washed 5 times with Hanks Balanced Salt Solution (HBBS) and incubated in 0.5% Trypsin-EDTA for 15 min. Cortices were then washed 5 times with MEM-HS supplemented with L-glutamine, essential amino acids, non-essential amino acids and 10% horse serum. The single cells in suspension were pelleted at 200 g for 5 min, and seeded at a density of 5 x 10^5^ cells/ml. Neural stem cells were cultured in DMEM-F12 with 5% FCS, B27 supplement and 20 ng/ml basic fibroblast growth factor (bFGF) on poly-D-lysine-coated culture dishes at 37°C in 5% CO_2_ (Kilpatrick & Bartlett, 1993; Johe *et al*, 1996; Azari *et al*, 2011; Mukhtar *et al*, 2020). Every second day, the culture medium was supplemented with 20 ng/ml bFGF. At 95% confluency cells were diluted 1:2. Differentiation was induced in neurobasal medium with B27 supplement/0.25x Glutamax five days after seeding.

For quantitative RT-PCR analysis or RNA-seq experiments, 3x10^5^ NSC were seeded in 2 ml medium on 35 mm dishes. For microscopy experiments, 1.6x10^5^ NSC were seeded in 1 ml medium on 12.8 mm dishes equipped with 12 mm coverslips.

Sh-mediated knockdown experiments were started one day after seeding by addition of 5 µl virus per 35 mm dish or 3 µl KD virus per 12.8 mm dish. To restore *RUS* expression, 10 µl or 6 µl *RUS* overexpression-virus per 35 mm or 12.8 mm dish, respectively, was added to KD cells four days after seeding.

### Cultivation of Neuro2A cells

Neuro2A cells were cultured in DMEM-Glutamax and 10% FCS at 37°C in 5% CO_2_.

### Immunohistochemistry

Cells were plated on poly-L-lysine-coated glass plates in a 24-well plate. All cell washes were done in PBS, all incubations were at RT. Cells were fixed in 4% paraformaldehyde (PFA) for 20 min at RT, washed once for 10 min and blocked with blocking solution (0.3% Triton X- 100, 2% donkey serum in PBS) for 30 min. The primary antibody (1:1000) was diluted in 200 µl blocking solution and added for 1.5 h while shaking. The antibody solution was removed, and the cells were washed three times for 10 min. Cells were incubated with the secondary antibody (1:2000) in 200 µl blocking solution for 1.5 h as before. After three 10-minute washes nuclei were stained for 15 min using DAPI (2-[4-Amidinophenyl]-6-indolecarbamidine dihydrochloride,) 1:1000 in PBS. The cells were mounted in the presence of diazabicyclo- octane (DABCO). Stained cells were analyzed with a Leica DM8000 fluorescent microscope, and images were quantitatively processed with ImageJ. Images from DAPI and antibody staining were thresholded, colocalized and watershed-transformed. The particles in the resulting overlay image were counted using the particle analyzer. Per experiment, 3-5 microscope fields on 3-4 plates each were recorded and analyzed.

### BrdU-labeling

Cull culture medium was supplemented with 1 µg/ml bromodesoxyuridine. After 24 hours, cells were immunostained with an anti-BrdU antibody.

### Quantitative reverse transcription-PCR (RT-qPCR)

RNA from cells, tissues or biochemical experiments was extracted with Trizol and chloroform and precipitated using 50% isopropanol and 15 µg linear acrylamide. RNA was washed twice with 75% EtOH, dissolved in nuclease-free water and reverse-transcribed using MMuLV RT (Thermo Fischer) and oligo(dT18-20). ChIRP and RIP-purified RNA was amplified with random hexamers. Quantitative RT-PCR analysis was performed with 1 µM of each primer in Fast SYBR® Green Master Mix (Thermo Fischer). The ΔCt values were normalized with amplicons detecting against TATA-binding protein (TBP) mRNA.

### 3’ RACE

The RUS 3’-end was cloned from a hippocampal RNA using the FirstChoice™ RLM-RACE Kit (Thermo Fischer). One microgram of RNA was reverse-transcribed using an anchored 3′ RACE oligo(dT) primer. This was followed by two rounds of nested PCR using RUS-3’-RACE as forward and 3’-outer primers and 3’-inner as reverse primer. The PCR product was gel-purified and sequenced.

### Generation of the RUS knockdown vector

ShRNAs were designed according to standard procedures (Yuan *et al*, 2004). In brief, 100 pmol RUS-sh-FW and 100 pmol RUS-sh-RV were annealed in 50 µl NEB2.1. The annealed fragment was cloned into pLKO.1-TRC-Puro vector, linearized with AgeI and EcoRI (Moffat *et al*, 2006) and amplified in Dh5α. For pLKO.1 vectors containing GFP as a selection marker, the puromycin resistance gene was replaced with the GFP gene via BamHI and KpnI restriction sites. Towards this end, the GFP cDNA was amplified from pLenti-CMV-GFP-Hygro (Campeau *et al*, 2009) by PCR using the primers: BamH-GFP-fw and Kpn-GFP-rv.

### Construction of pcDNA-5FRT-5xMS2

pcDNA.5-FRT vectors used to generate stable FlpIN Neuro2 A cells were equipped with 5xMS2 stem-loops. The 3xMS2 stem-loop sequence was PCR-amplified with the primers MS2_fw and MS2_rv from pAdMl3-(MS2)_3_, digested with BamHI and XbaI, and ligated to BamHI/XbaI-linearized pcDNA5-FRT. Upon amplification in Dh5α, one clone fortuitously expanded 3xMS2 stem-loops to 5xMS2 stem-loops. This clone was used.

### Generation of RUS overexpression vector

*RUS* and Δ5’*RUS* sequences were isolated from a hippocampal cDNA library by PCR with the primers: RUS-LIC-fw or Δ5’ RUS-LIC-fw, respectively and RUS-LIC-rv and cloned into pcDNA-5-FRT or pcDNA.5-FRT-5xMS2 (Thermo Fischer) via LIC cloning (Wang *et al*, 2012) and amplified in Dh5α. The *RUS* cDNA was the shuffled into pLenti-CMV-GFP-Hygro (Campeau *et al*, 2009) via ClaI and ApaI restriction sites to replace GFP and the hygromycin resistance gene. All constructs were verified by sequencing.

### Construction of pLenti-FRT

pLenti-GFP-Puro (Campeau *et al*, 2009) was digested with XbaI and BamHI to remove GFP downstream of the CMV promoter. FRT site was generated by annealing the oligonucleotides FRT_fw and FRT_rv. For annealing, 100 pmol of each oligonucleotide was heated in 50 µl NEB 2.1 to 95°C for 5min and slowly cooled down. 2 µl annealing scale was ligated into 20 ng digested vector and transformed in Dh5α.

### Production of lentiviral particles

All lentiviral experiments were conducted according to standard protocols (Moffat *et al*, 2006) and approved by the Bavarian state. 3x10^6^ HEK293T cells were seeded in 8 ml DMEM- GlutMax supplemented with 8% FCS on a 10 cm culture dish. Per virus production, 4 10 cm dishes were seeded. Next day, 53 µg DNA in a molar ratio of 2:1:1 of lentiviral-vector:psPAX2:pMD2.G transfected into 50-70% confluent cells. The medium was changed next day. Two days after transfection, viral particles were purified by sedimentation (87.000 g, 2h) from the medium and dissolved in 200 µl TBS5 (50 mM Tris-HCl pH 7.8, 130 mM NaCl, 10 mM KCl, 5 mM MgCl_2_, 10% BSA).

### Subcellular fractionation

Subcellular fractionation was adapted from (Gagnon *et al*, 2014). Briefly, cells were lysed in ice-cold hypotonic lysis buffer (HLB; 10 mM Tris-HCl pH 7.5, 10 mM NaCl, 3 mM MgCl_2_, 0.3% NP-40, 10% glycerol) for 10 min on ice. The cytoplasm was harvested by centrifugation (1000 g, 5 min) and the nuclear pellet was washed thrice in HLB. Nuclei were incubated in ice-cold Modified Wuarin-Schibler buffer (MWS; 10 mM Tris-HCl pH 7.5, 4 mM EDTA, 0.3 M NaCl, 1 M urea, 1% NP-40) for 15 min on ice. The nucleoplasm was separated from the chromatin by centrifugation (1000 g, 5 min). The RNA in the cytoplasmic and nucleoplasmic fractions was ethanol-precipitated and subjected along with the chromatin pellet for RNA purification.

### RNA-seq analysis

Total RNA was isolated and polyA-enriched. After reverse transcription, the cDNA was fragmented, end-repaired, and polyA-tailed. Solexa sequencing adaptors were ligated, and adaptor-modified fragments were enriched by 10-18 cycles of PCR amplification. Quantity and the size of the sequencing library were accessed on a Bioanalyzer before sequencing on an Illumina NextSeq 500 platform. Sequencing reads from FASTAQ files were aligned the STAR Aligner version (Dobin *et al*, 2013) and quantified using rsem (Li & Dewey, 2011). The reference genome used for alignment was constructed using the mm10 fasta file and GRCm38.99 transcript table. Quantified values were further statistically evaluated using Bioconductor’s DeSeq2 package (Love *et al*, 2014). Expression changes with an FDR < 0.05 were considered significant. Among them, genes with a stat < -2 or >2 were extracted as down- or upregulated genes, respectively (Table EV2).

### Gene ontology term enrichment analysis

GO enrichment employed the web-based PANTHER software (Mi *et al*, 2013). The deregulated genes enriching for GO terms of interest were extracted from the provided xml file and matched to their expression values using R.

### ChIRP-seq analysis

NSCs from 8 x 15 cm dishes were harvested 7 days after seeding and washed twice with PBS. Cells were crosslinked in 100 ml 1% glutaraldehyde for 10 min at RT. Crosslinking was quenched 125 mM glycine for 5 min. Cells were pelleted at 1000 g for 5 min. ChIRP was done according to (Chu *et al*, 2011). Crosslinked cells were washed twice in PBS and lysed in 2 ml ChIRP-lysis buffer (50 mM Tris-HCl pH 7.0, 10 mM EDTA, 1% SDS, 1 mM PMSF, 1x protease inhibitor, SuperaseIn 100 U/ml). Chromatin shearing by Bioruptor typically yielded fragments of 150-600 bp. Sheared chromatin was diluted with 4 ml ChIRP-hybridization buffer (50 mM Tris-HCl pH 7.0, 750 mM NaCl, 15% (m/v) formamide, 1 mM EDTA, 1% SDS, with protease and RNase inhibitors) and divided into two aliquots, which were hybridized with 100 pmol biotinylated ‘odd’ and ‘even’ probe sets, respectively, at 37°C for 4 h with continuous rotation. Then 1 mg of magnetic streptavidin bead suspension (Thermo Fischer) in ChIRP-Lysis buffer were added and incubated for 30 min at 37°C with continuous rotation. Beads were washed 5 times with 1 ml ChIRP Wash buffer (300 mM NaCl, 30 mM Na_3_-citrate, 0.1% SDS, 1 mM PMSF) for 5 min at 37°C. 90% of bead material was used for DNA isolation and 10% for RNA isolation. The enrichment of *RUS*, TBP mRNA, *MALAT*, and *XIST* was analyzed by RT-qPCR.

Isolated DNA was processed alongside an input chromatin sample. Ends were blunted with T4 DNA polymerase and polynucleotide kinase and an AMP was added. Solexa sequencing adaptors were ligated and adaptor-modified fragments were enriched by 10-18 cycles of PCR amplification. Sequencing libraries were size-selected on AMPure Beads (Beckman Coulter), quality-controlled on a Bioanalyzer (Agilent) and sequenced on an Illumina NextSeq-500 platform.

Sequencing reads from FASTQ files were aligned with bowtie2 (Langmead & Salzberg, 2012) to mm10. Multimapping reads were removed using samtools (Li *et al*, 2009). ChIRP peaks were called with MACS1.4 for both probe sets independently (Feng *et al*, 2012). The deeptools package was used to generate the bedgaph files (Ramírez *et al*, 2016). Bedtools (Quinlan & Hall, 2010) and python 2.7 matched even and odd bedgraph files into a single bedgraph file via the ’take-lower’ method. The experiment was done in triplicates Only peaks occurring in each even and odd sample and in all three data sets called with Bioconductor’s GenomicRanges package (Lawrence *et al*, 2013) were considered valid *RUS* binding sites. The overlap demanded a minimal distance of 200 bp between the ‘even’ and ‘odd’ summit. Probe sequences within overlapping peaks were detected using Fimo (Grant *et al*, 2011) of the MEME software (Bailey *et al*, 2015) and removed using a cutoff of p <1e-8 before further analysis using GenomicRanges (Lawrence *et al*, 2013).

Filtered peaks were annotated with Homer (Heinz *et al*, 2010) using mm10 as reference genome (Table EV3). The obtained annotation statistic was used to calculate the distribution of *RUS* peaks within promoter, intergenic, intron, and close to repetitive sites. The annotated neighboring genes of *RUS* peaks were considered putative *RUS* target genes. GO term enrichment of putative target genes employed the web-based PANTHER software (Mi et al, 2013). Next, putative target gene expression and changes upon in shRNA*^CON^*and shRNA*^RUS^* treatment on day 5 and 7 were extracted from the RNA-seq data using the R-package SummarizedExperiments and DeSeq2 (Table EV3). Expression changes with an FDR < 0.05 were considered significant. Among them, genes with a stat < -2 or >2 were considered as down- or upregulated genes. Expression values of both time points were merged, log2- transformed and ranked by hierarchically clustering using the Euclidean Distance method in R. Furthermore, the correlation between *RUS* and putative target gene expression was calculated using the Pearson correlation coefficient on both time points separately (Table EV3).

### MS2 Affinity purification of RUS interactors

Stable pools of Neuro2A cells expressing 5xMS2-tagged *RUS* were generated as follows. 5x10^4^ Neuro2A cells were transfected with 5 µl pLenti-FRT virus and 2 days later selected in GlutMax, 8% FCS supplemented with 2 µg/ml puromycin and expanded. 10^6^ Neuro2A-FRT cells were seeded on a 10 cm culture dish. On the next day, cells were transfected with 15 µg plasmid DNA, consisting of a molar ratio of 1:6 (up to 1:9) of pcDNA5-lncRNA-5xMS2: pCSFLPe (encoding the flipase). Plasmids were diluted appropriately in 300 µl 150 mM NaCl and 15 µl JetPEI (2.6 µg/µl) and mixed. After 30 min equilibration at RT, the solution was added dropwise to Neuro2-FRT cells. Two days later, cells were transferred to a new 10 cm dish and selected in GlutMax 8% FCS, 2 µg/ml puromycin, and 600 µg/ml hygromycin. The medium was replaced every second day to remove cell debris. Colonies formed 7-10 days after transfection. They were harvested and further cultivated.

Nuclear extract from MS2-tagged *RUS*-expressing Neuro2A cells was prepared typically from 8x10^7^ cells without dialysis, according to (Dignam *et al*, 1983). Extract preparation and MS2-affinity purification were done at 4°C. Cell pellets were suspended in 5 vol buffer A (10 mM HEPES pH 7.9 at 4°C, 1.5 mM MgCl_2_, 10 mM KCl, 0.5 mM DTT, 200 U/ml RNAsin) and incubated for 10 min. Cells were homogenized with a Dounce tissue grinder. Nuclei were pelleted at 500 g for 10 min, washed with 5 nuclear volumes (vol) buffer A, dissolved in one vol buffer C (20 mM HEPES pH 7.9, 25% (v/v) glycerol, 0.42 M KCl, 1.5 mM MgCl_2_, 0.2 mM EDTA, 0.5 mM PMSF, 0.5 mM DTT, 200 U/ml RNAsin) and homogenized again with a Dounce tissue grinder. After gentle rotation for 30 min, chromatin was pelleted at 17.000 g for 30 min. The supernatant was diluted with 1 vol buffer G (20 mM HEPES pH 7.9, 20% (v/v) glycerol, 0.2 mM EDTA, 0.5 mM PMSF, 0.5 mM DTT, 200 U/ml RNAsin) and used for affinity purification.

Standard MS2-affinity purification was done on supernatant containing 1 mg protein. To this, 760 pmol yeast t-RNA competitor and 120 pmol recombinant MS2BP-MBP (Jurica *et al*, 2002; Zhou & Reed, 2003) was added. After 2 h of gentle rotation, 50 µl equilibrated amylose resin (New England Biolabs) was added and incubation continued for 2 h. The resin was pelleted at 1900 g for 1 min and washed thrice with 900 µl buffer D (buffer G containing 0.1 M KCl and lacking RNasin) and thrice 900 µl buffer F (buffer D containing 1.5 mM MgCl_2_,).

RNA-interacting proteins were identified by mass spectrometry. Interacting proteins were eluted with 50 µg RNAse A in 80 µl buffer D at 37°C for 10 min. The resin was pelleted at 1900 g for 1 min at 4°C and the supernatant subjected to Filter Aided Sample Preparation (Wiśniewski *et al*, 2009) and peptides were desalted using C18 StageTips, dried by vacuum centrifugation and dissolved in 20 µL 0.1% formic acid. Samples were analyzed on a Easy nLC 1000 coupled online to a Q-Exactive mass spectrometer (Thermo Scientific, US). Eight µl peptide solution per sample were separated on a self-packed C18 column (30 cm × 75 µm; ReproSil-Pur 120 C18-AQ, 1.9 μm, Dr. Maisch GmbH, Germany) using a 180 min binary gradient of water and acetonitrile supplemented with 0.1% formic acid (0 min., 2% B; 3:30 min., 5% B; 137:30 min., 25% B; 168:30 min., 35% B; 182:30 min., 60% B) at 50 °C column temperature. A top 10 DDA method was used. Full scan MS spectra were acquired with a resolution of 70,000. Fragment ion spectra were recorded using a 2 m/z isolation window, 75 ms maximum trapping time with an AGC target of 10^5^ ions.

The Raw Data were analyzed with the MaxQuant (version 2.0.1.0) software (Cox & Mann, 2008) using a one protein per gene canonical database of Mus musculus from Uniprot (download : 2021-04-09; 21998 entries). Trypsin was defined as protease. Two missed cleavages were allowed for the database search. The option first search was used to recalibrate the peptide masses within a window of 20 ppm. For the main search peptide and peptide fragment mass tolerances were set to 4.5 and 20 ppm, respectively. Carbamidomethylation of cysteine was defined as a static modification. Acetylation of the protein N-terminus as well as oxidation of methionine set as variable modifications. Match between runs was enabled with a retention time window of 1 min. Two ratio counts of unique peptides were required for label-free quantification (LFQ).

Output files were further analyzed using the software Perseus (Tyanova *et al*, 2016). Proteins identified by site, reverse matching peptides and contaminants were removed and LFQ intensities were log2 transformed. Next, only protein groups with 5 out of 5 quantifications in one condition were considered for relative protein quantification. To account for proteins that were only consistently quantified in one condition, data imputation was used with a down-shift of 2 and a width of 0.2. A permuation based FDR correction (Tusher *et al*, 2001) for multiple hypotheses was applied (p = 0.05; s0 = 0.1). Proteins were considered enriched if the fold change was greater than two and the p-value less than 0.002 (Table EV4).

### RNA immunoprecipitation

Protein A/G-Agarose beads (35 µl, Thermo Fischer) were blocked overnight with 1% BSA in buffer D. To nuclear extract from 5x10^6^ NSC 760 pmol yeast t-RNA, 300 µg salmon sperm DNA and 4 µg Lbr antibody was added and incubated for 2 h at 4°C under gentle rotation. Anti-rabbit IgG was used as a negative control. The binding reaction was added to blocked Protein A/G-Agarose and incubated for 2 h at 4°C with gentle rotation. Protein A/G beads were sedimented, washed 5x with 900 µl buffer D, suspended in 800 µl Trizol and subject to RNA extraction. RUS levels were analyzed by RT-PCR analysis and compared against the IgG purification. The experiment was performed in triplicates and statistically evaluated by a one-tailored student’s t-test using Bonferroni p-value adjustment.

## Structured Methods

## Acknowledgements

We thank Aline Campos, Silke Krause and Anna Berghofer for technical assistance, Tobias Straub for advice on bioinformatic analysis, Bianka Baying, Vladimir Benes (EMBL GeneCore), and Stefan Krebs (LAFUGA) for library preparation and sequencing and Christian Haass for his continued support. We are grateful to Sandra Schick, Marie Kube and Rodrigo Villaseñor for critical reading of the manuscript. This work was funded by the Deutsche Forschungsgemeinschaft (DFG) within the framework of the Munich Cluster for Systems Neurology (EXC 2145 SyNergy– ID 390857198), grant Be1140/8-1 (to PBB) and the Adele Hartmann Programm of the LMU (to JCS).

## Author contributions

JCS and MS conceived the study. MS and VM performed experiments. MS analyzed and interpreted all experiments. SAM and SFL performed LC-MS analysis. MS and PBB wrote the manuscript. All authors proofread the manuscript.

## Data Availability Section

The datasets and computer code produced in this study are available in the following databases:

- RNA-Seq data: Gene Expression Omnibus GSE196487 (https://www.ncbi.nlm.nih.gov/geo/query/acc.cgi?acc=GSE196487, reviewer access token : szajouyirxsjfqj)
- ChiRP-Seq data: Gene Expression Omnibus GSE196527 (https://www.ncbi.nlm.nih.gov/geo/query/acc.cgi?acc=GSE196527, reviewer access token: kbytkaqghfgfzuf)
- Protein interaction AP-MS data: PRIDE PXD031664 (Perez-Riverol *et al*, 2022) (http://www.ebi.ac.uk/pride/archive/projects/PXD000208, reviewer access: username, password: 6p26fYpI)
- Computer scripts: GitHub (https://github.com/MariusFSchneider/Schneider22)

## Tables and their legends

**Table EV1:** Used oligonucleotides

**Table EV2:** Differential gene expression analysis measured by RNASeq

**Table EV3:** RUS genomic binding sites measured by ChIRP-Seq and expression of putative target genes

**Table EV4:** Statistical evaluation of identified and quantified proteins after RUS affinity purifications by LC-MS.

## Expanded View Figure legends

**Figure EV1.**
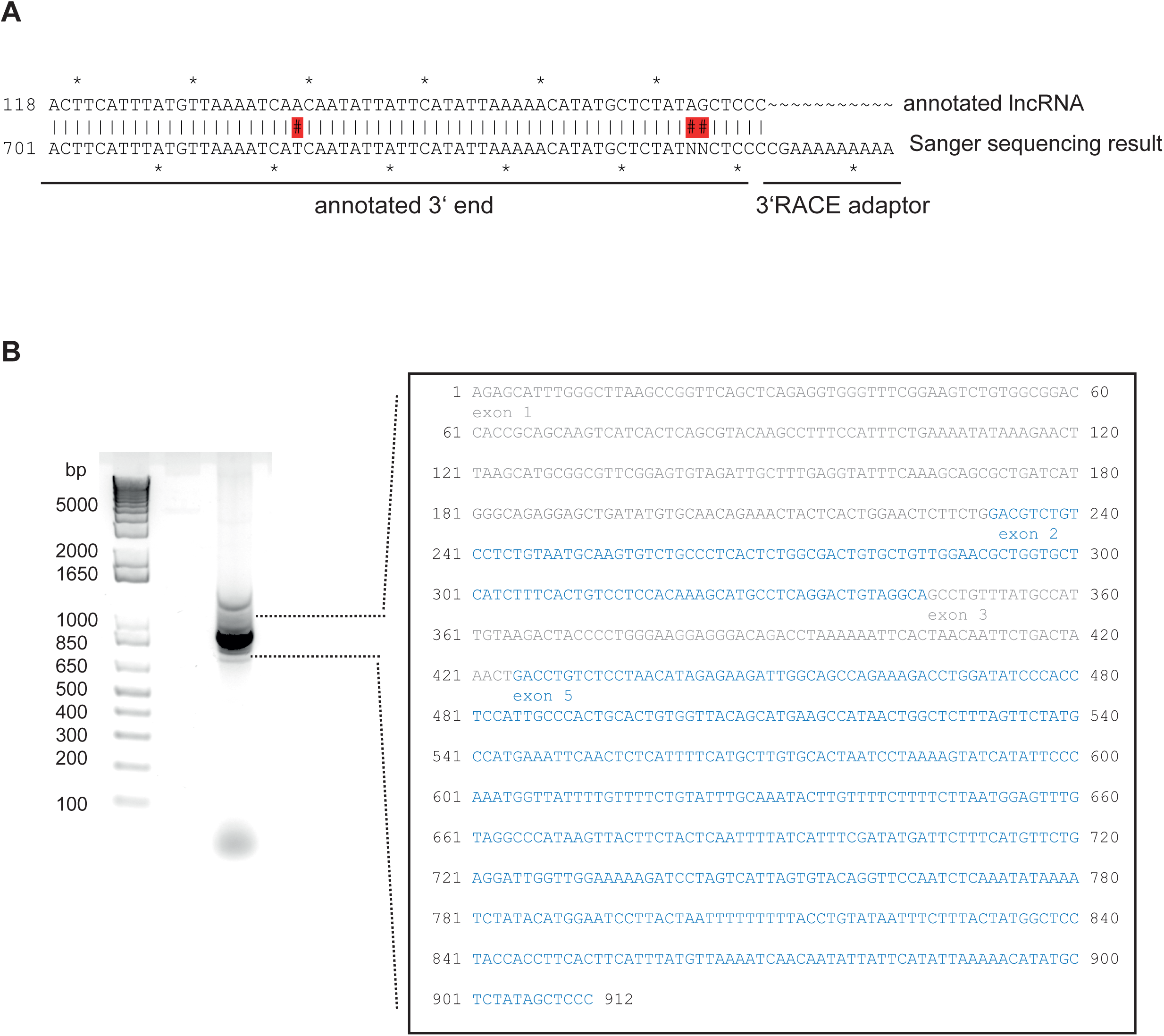
Molecular characterization of RUS. **A.** Sequence of the *RUS* 3’ end determined by 3’ RACE. **B.** Left: RT-PCR amplification of *RUS* using primers annealing to 5’ and 3’ ends, revealing the dominant isoform *RUS-1*. Right panel: Sequence of *RUS-1* (912 nt). Note that the annotated exon 4 is missing.

**Figure EV2.**
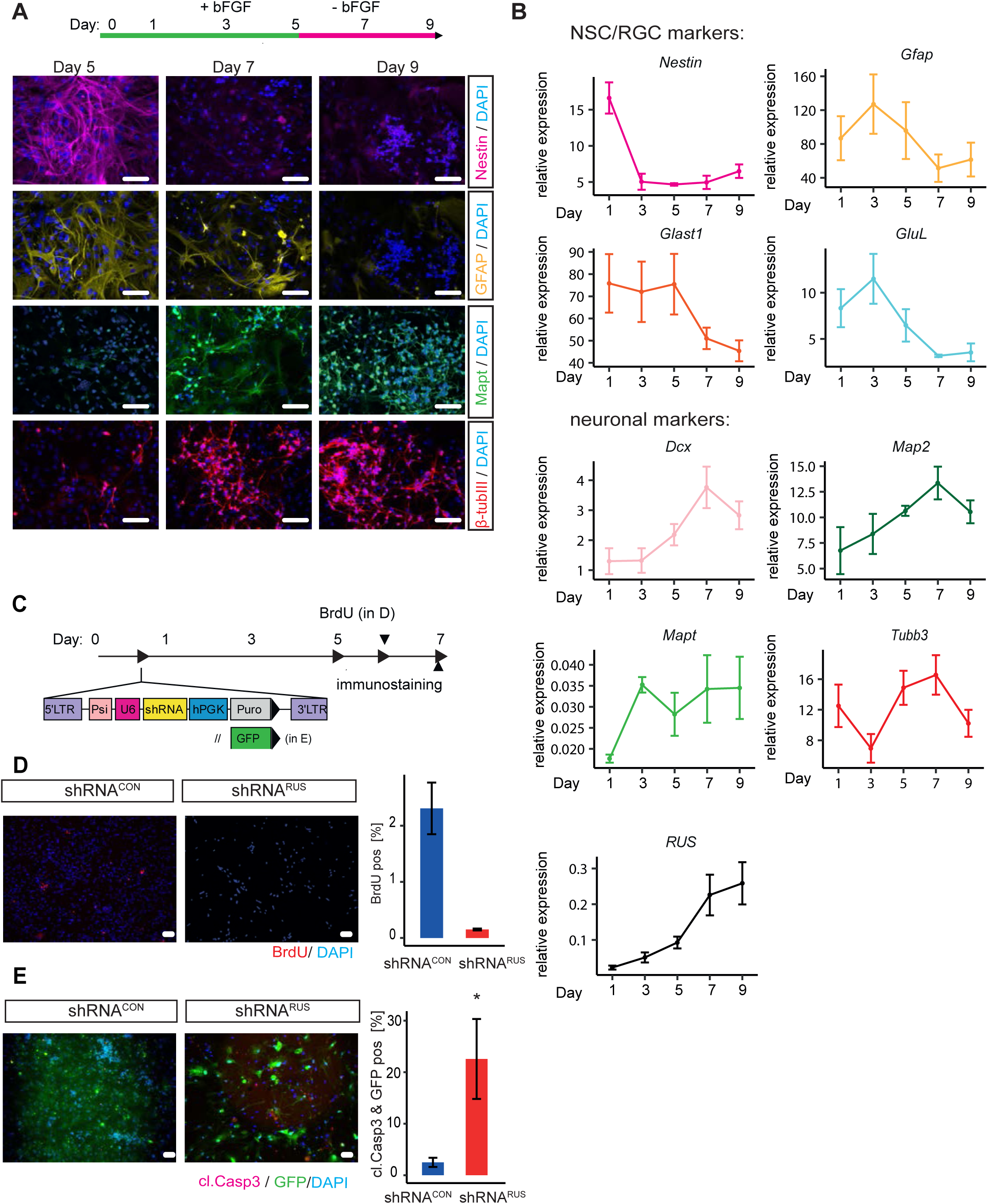
Cell loss upon *RUS* depletion through reduced cell proliferation and increased apoptosis. A. Immunostaining of differentiating embryonic cortical NSC for the NSC marker nestin (magenta, top row), the RGC marker Gfap (yellow, 2^nd^ row), and for the neuronal markers Mapt (green, 3^rd^ row) and β-tubulin III (red, bottom row) on the indicated days. Cell nuclei stained with 4′,6-Diamidin-2-phenylindol (DAPI), bar = 25 µm. B. Quantitative RT-PCR analysis of mRNA encoding Dcx (double cortin), Gfap (glial fibrillary acidic protein), the glutamate transporter Glast, glutamate-ammonia ligase (GluL), Map2 (microtubule-associated protein 2), Mapt (microtubule-associated protein tau), nestin and β-tubulin III during differentiation of embryonic cortical NSC. Values are relative to the constitutively expressed TATA-binding protein (TBP) mRNA values in the same preparations, which were also used for normalization. Error bars: standard error of the mean of 3 independent experiments. C. Experimental strategy to deplete *RUS* in differentiating NSC using lentiviral shRNAs. LTR: long terminal repeat, psi: packing signal, U6: U6-promoter, hPKG: human phosphoglycerate kinase promoter, Puro: puromycin resistance gene, GFP: Green fluorescent protein. D. Quantification of BrdU immunostaining as a measure of replication. BrdU was added to differentiating NSC cultures on day 6 and its incorporation measured by immunostaining in control (shRNA*^CON^*) and knockdown (shRNA*^RUS^*) cells using specific antibodies (red). Nuclei were stained with DAPI. Scale bar = 25 µm. Images were quantified with ImageJ (right panel). Error bars show the standard deviation of three independent experiments (* p < 0.05, ** p < 0.01, *** p < 0.005). E. Quantification of cleaved Caspase-3 immunostaining as a measure of apoptosis. Cleaved Caspase-3 was detected in GFP-expressing control (shRNA*^CON^*) and knockdown (shRNA*^RUS^*) cells using specific antibodies (magenta). Nuclei were stained with DAPI. Scale bar = 25 µm. The stainings were quantified with ImageJ (right panel). Error bars show the standard deviation of three independent experiments (* p < 0.05, ** p < 0.01, *** p < 0.005).

**Figure EV3.**
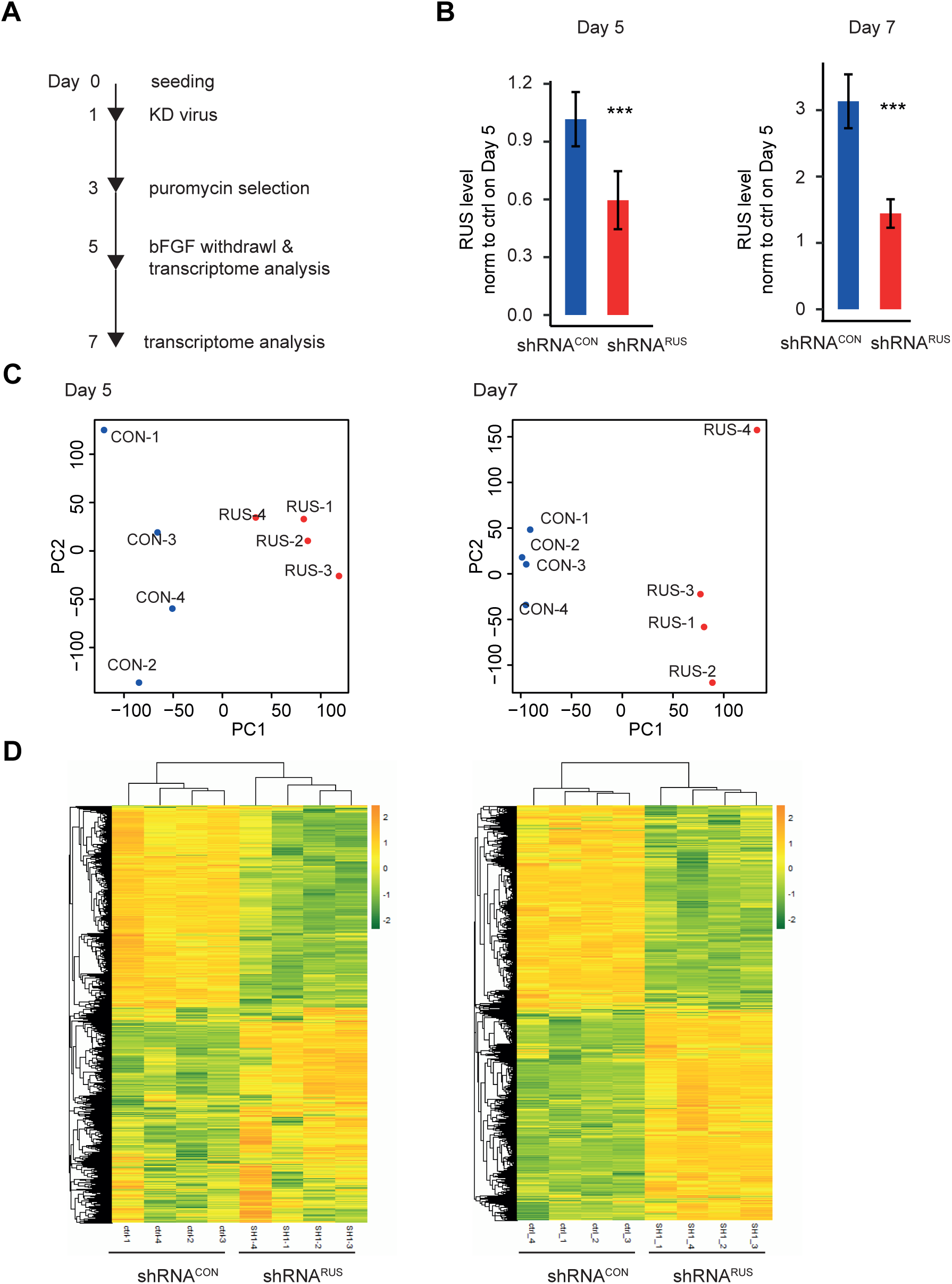
Transcriptome changes upon depletion of RUS. **A.** Experimental strategy and timeline for transcriptome analysis. **B.** *RUS* levels determined by RT-qPCR in *RUS* knockdown cells (red, expressing shRNA*^RUS^*) compared to control cells (blue, expressing a scrambled shRNA*^CON^*). Error bars show the standard deviation of the mean of four experiments. **C.** Principle component analysis comparing RNA-seq profiles of cells treated with shRNA*^CON^*(*CON*) or shRNA*^RUS^*(*RUS*) on day 5 (left) or day 7 (right). The four replicates are labeled 1-4. **D.** Heatmap displaying the relative expression of genes with significant deregulation (FDR < 0.05) on days five (left) and seven (right), hierarchically clustered using the Euclidean distance.

**Figure EV4.**
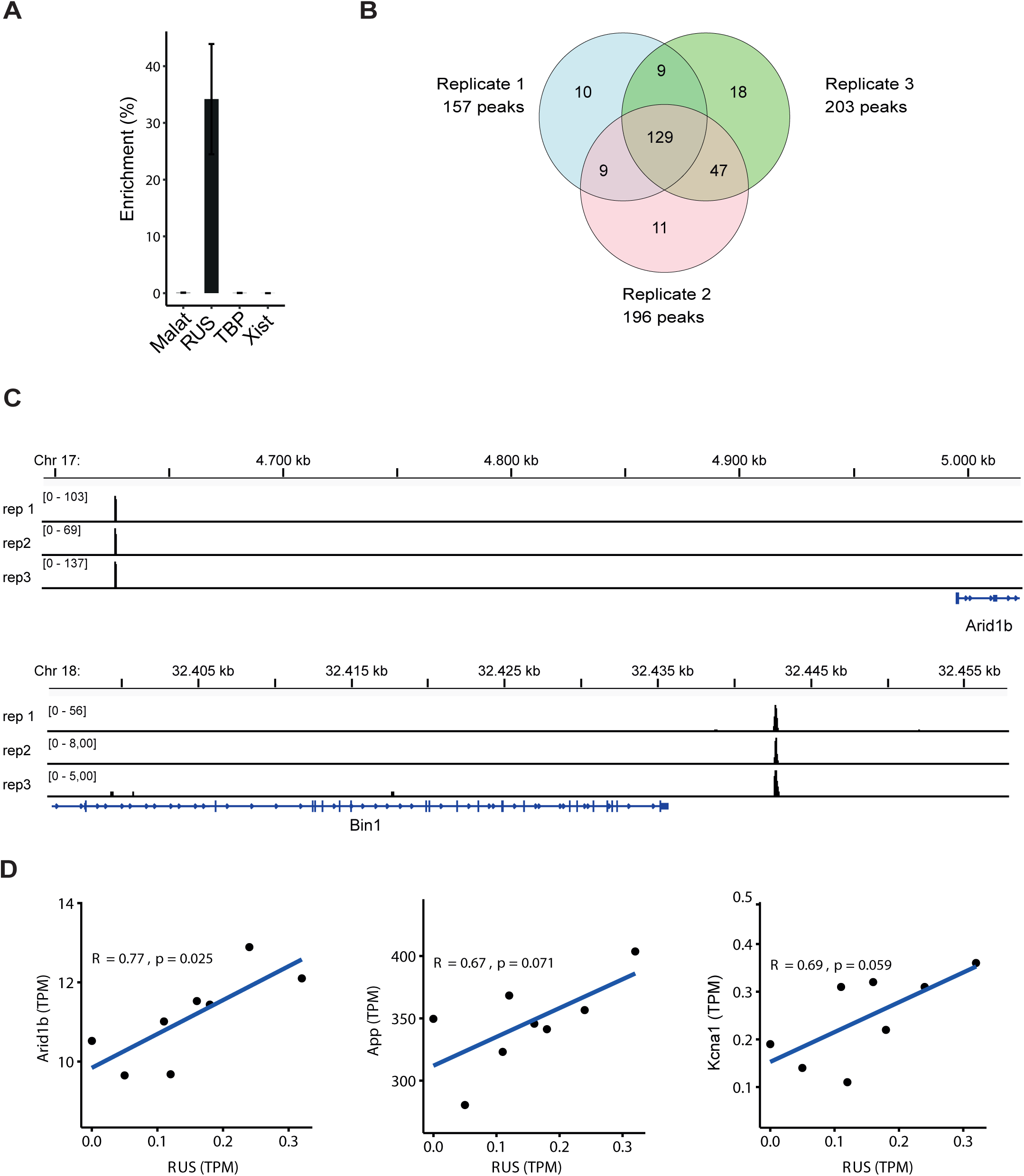
Chromatin localization of RUS determined by ChIRP. **A.** Enrichment of *RUS*, *MALAT*, *TBP* mRNA, and *XIST* in ChIRP from differentiated NSC expressed as percent of the value obtained from input chromatin. Error bars: the standard deviation of 3 experiments. **B.** Venn diagram showing the number of ChIRP peaks obtained in the three independent experiments and their overlap. **C.** Browser view of two examples of *RUS* localization close to relevant neurogenic genes. The *RUS* ChIRP tag density of the three replicates is plotted in separate tracks in the genomic regions of the *Arid1b* (top) and *Bin1* (bottom) genes. For orientation, the respective chromosomal regions are displayed above and the gene models below the traces. **D.** Correlation analyses showing the relationship between *RUS* expression and three selected putative target genes of cluster II, *App*, *Arid1* and *Kcna1,* in control (shRNA*^CON^*) and *RUS* knockdown (shRNA*^RUS^*) cells on day 5.

## References

Andrews BJ, Proteau GA, Beatty LG & Sadowski PD (1985) The FLP recombinase of the 2μ circle DNA of yeast: Interaction with its target sequences. Cell 40: 795–803

Antonelli F, Casciati A, Tanori M, Tanno B, Linares-Vidal M V, Serra N, Bellés M, Pannicelli A, Saran A & Pazzaglia S (2018) Alterations in Morphology and Adult Neurogenesis in the Dentate Gyrus of Patched1 Heterozygous Mice. Front Mol Neurosci 11: 168

Aruga J, Yokota N & Mikoshiba K (2003) Human SLITRK family genes: Genomic organization and expression profiling in normal brain and brain tumor tissue. Gene 315: 87–94

Azari H, Osborne GW, Yasuda T, Golmohammadi MG, Rahman M, Deleyrolle LP, Esfandiari E, Adams DJ, Scheffler B, Steindler DA, et al (2011) Purification of immature neuronal cells from neural stem cell progeny. PLoS One 6

Bailey TL, Johnson J, Grant CE & Noble WS (2015) The MEME Suite. Nucleic Acids Res 43: W39– W49

Briggs JA, Wolvetang EJ, Mattick JS, Rinn JL & Barry G (2015) Mechanisms of Long Non-coding RNAs in Mammalian Nervous System Development, Plasticity, Disease, and Evolution. Neuron 88: 861–877

Campeau E, Ruhl VE, Rodier F, Smith CL, Rahmberg BL, Fuss JO, Campisi J, Yaswen P, Cooper PK & Kaufman PD (2009) A Versatile Viral System for Expression and Depletion of Proteins in Mammalian Cells. PLoS One 4: e6529

Ceballos-Chávez M, Rivero S, García-Gutiérrez P, Rodríguez-Paredes M, García-Domínguez M, Bhattacharya S & Reyes JC (2012) Control of neuronal differentiation by sumoylation of BRAF35, a subunit of the LSD1-CoREST histone demethylase complex. Proc Natl Acad Sci U S A 109: 8085–8090

Chen J, Dong X, Cheng X, Zhu Q, Zhang J, Li Q, Huang X, Wang M, Li L, Guo W, et al (2021) Ogt controls neural stem/progenitor cell pool and adult neurogenesis through modulating Notch signaling. Cell Rep 34: 108905

Chou S-M, Li K-X, Huang M-Y, Chen C, King Y-HL, Li GG, Zhou W, Teo CF, Jan YN, Jan LY, et al (2021) Kv1.1 channels regulate early postnatal neurogenesis in mouse hippocampus via the (TrkB) signaling pathway. Elife 10: e58779

Chu C, Qu K, Zhong FL, Artandi SE & Chang HY (2011) Genomic maps of long noncoding RNA occupancy reveal principles of RNA-chromatin interactions. Mol Cell 44: 667–78

Chun-Kan C, Mario B, Constanza J, Erik A, Noah O, Christine S, Amy C, Andrea C, Patrick M & Mitchell G (2016) Xist recruits the X chromosome to the nuclear lamina to enable chromosome-wide silencing. Science (80-) 354: 468–472

Clark BS & Blackshaw S (2017) Understanding the Role of lncRNAs in Nervous System Development. In Long Non Coding RNA Biology pp 253–282. Singapour: Springer Nature Singapore Pte Ltd

Cox J & Mann M (2008) MaxQuant enables high peptide identification rates, individualized p.p.b.- range mass accuracies and proteome-wide protein quantification. Nat Biotechnol 26: 1367–1372

Cox J & Mann M (2009) Computational principles of determining and improving mass precision and accuracy for proteome measurements in an Orbitrap. J Am Soc Mass Spectrom 20: 1477–85

Dignam JD, Lebovitz RM & Roeder RG (1983) Accurate transcription initiation by RNA polymerase II in a soluble extract from isolated mammalian nuclei. Nucleic Acids Res 11: 1475–1489

Djebali S, Davis C a., Merkel A, Dobin A, Lassmann T, Mortazavi A, Tanzer A, Lagarde J, Lin W, Schlesinger F, et al (2012) Landscape of transcription in human cells. Nature 489: 101–108

Dobin A, Davis CA, Schlesinger F, Drenkow J, Zaleski C, Jha S, Batut P, Chaisson M & Gingeras TR (2013) STAR: Ultrafast universal RNA-seq aligner. Bioinformatics 29: 15–21

Engler A, Rolando C, Giachino C, Saotome I, Erni A, Brien C, Zhang R, Zimber-Strobl U, Radtke F, Artavanis-Tsakonas S, et al (2018) Notch2 Signaling Maintains NSC Quiescence in the Murine Ventricular-Subventricular Zone. Cell Rep 22: 992–1002

Engreitz JM, Ollikainen N & Guttman M (2016) Long non-coding RNAs: Spatial amplifiers that control nuclear structure and gene expression. Nat Rev Mol Cell Biol 17: 756–770

Eriksson PS, Perfilieva E, Björk-Eriksson T, Alborn A-M, Nordborg C, Peterson DA & Gage FH (1998) Neurogenesis in the adult human hippocampus. Nat Med 4: 1313–1317

Feng J, Liu T, Qin B, Zhang Y & Liu XS (2012) Identifying ChIP-seq enrichment using MACS. Nat Protoc 7: 1728–1740

Franks TM, McCloskey A, Shokirev M, Benner C, Rathore A & Hetzer MW (2017) Nup98 recruits the Wdr82-Set1A/COMPASS complex to promoters to regulate H3K4 trimethylation in hematopoietic progenitor cells. Genes Dev 31: 2222–2234

Fusaro G, Dasgupta P, Rastogi S, Joshi B & Chellappan S (2003) Prohibitin Induces the Transcriptional Activity of p53 and Is Exported from the Nucleus upon Apoptotic Signaling. J Biol Chem 278: 47853–47861

Gagnon KT, Li L, Janowski BA & Corey DR (2014) Analysis of nuclear RNA interference in human cells by subcellular fractionation and Argonaute loading. Nat Protoc 9: 2045–2060

Galichet C, Guillemot F & Parras CM (2008) Neurogenin 2 has an essential role in development of the dentate gyrus. Development 135: 2031–2041

Garay PM, Wallner MA & Iwase S (2016) Yin–yang actions of histone methylation regulatory complexes in the brain. Epigenomics 8: 1689–1708

Gozalo A, Duke A, Lan Y, Pascual-Garcia P, Talamas JA, Nguyen SC, Shah PP, Jain R, Joyce EF & Capelson M (2020) Core Components of the Nuclear Pore Bind Distinct States of Chromatin and Contribute to Polycomb Repression. Mol Cell 77: 67–81.e7

Grant CE, Bailey TL & Noble WS (2011) FIMO: Scanning for occurrences of a given motif. Bioinformatics 27: 1017–1018

Hatakeyama J & Kageyama R (2006) Notch1 Expression Is Spatiotemporally Correlated with Neurogenesis and Negatively Regulated by Notch1-Independent Hes Genes in the Developing Nervous System. Cereb Cortex 16: i132--i137

Heinz S, Benner C, Spann N, Bertolino E, Lin YC, Laslo P, Cheng JX, Murre C, Singh H & Glass CK (2010) Simple Combinations of Lineage-Determining Transcription Factors Prime cis-Regulatory Elements Required for Macrophage and B Cell Identities. Mol Cell 38: 576–589

Hezroni H, Perry RBT & Ulitsky I (2019) Long noncoding RNAs in development and regeneration of the neural lineage. Cold Spring Harb Symp Quant Biol 84: 165–177

Hublitz P, Kunowska N, Mayer UP, Müller JM, Heyne K, Yin N, Fritzsche C, Poli C, Miguet L, Schupp IW, et al (2005) NIR is a novel INHAT repressor that modulates the transcriptional activity of p53. Genes Dev 19: 2912–2924

Imura T, Kornblum HI & Sofroniew M V. (2003) The predominant neural stem cell isolated from postnatal and adult forebrain but not early embryonic forebrain expresses GFAP. J Neurosci 23: 2824–2832

Ivanov D (2019) Notch Signaling-Induced Oscillatory Gene Expression May Drive Neurogenesis in the Developing Retina. Front Mol Neurosci 12: 226

Johansson HE, Liljas L & Uhlenbeck OC (1997) RNA recognition by the MS2 phage coat protein.Semin Virol 8: 176–185

Johe KK, Hazel TG, Muller T, Dugich-Djordjevic MM & McKay RDG (1996) Single factors direct the differentiation of stem cells from the fetal and adult central nervous system. Genes Dev 10: 3129–3140

Jurica MS, Licklider LJ, Gygi SP, Grigorieff N & Moore MJ (2002) Purification and characterization of native spliceosomes suitable for three-dimensional structural analysis. Rna 8: 426–439

Kilpatrick TJ & Bartlett PF (1993) Cloning and growth of multipotential neural precursors: Requirements for proliferation and differentiation. Neuron 10: 255–265

Kind J, Pagie L, Ortabozkoyun H, Boyle S, De Vries SS, Janssen H, Amendola M, Nolen LD, Bickmore WA & Van Steensel B (2013) Single-cell dynamics of genome-nuclear lamina interactions. Cell 153: 178–192

Kopp F & Mendell JT (2018) Functional Classification and Experimental Dissection of Long Noncoding RNAs. Cell 172: 393–407

Koushyar S, Jiang WG & Dart DA (2015) Unveiling the potential of prohibitin in cancer. Cancer Lett 369: 316–322

Langmead B & Salzberg SL (2012) Fast gapped-read alignment with Bowtie 2. Nat Methods 9: 357– 359

Lawrence M, Huber W, Pagès H, Aboyoun P, Carlson M, Gentleman R, Morgan MT & Carey VJ (2013) Software for Computing and Annotating Genomic Ranges. PLoS Comput Biol 9: 1–10

Li B & Dewey CN (2011) RSEM: accurate transcript quantification from RNA-Seq data with or without a reference genome. BMC Bioinformatics 12: 323

Li H, Handsaker B, Wysoker A, Fennell T, Ruan J, Homer N, Marth G, Abecasis G & Durbin R (2009) The Sequence Alignment/Map format and SAMtools. Bioinformatics 25: 2078–2079

Lin N, Chang KY, Li Z, Gates K, Rana Z, Dang J, Zhang D, Han T, Yang CS, Cunningham TJ, et al (2014) An evolutionarily conserved long noncoding RNA TUNA controls pluripotency and neural lineage commitment. Mol Cell 53: 1005–1019

Liu Z, Yan M, Liang Y, Liu M, Zhang K, Shao D, Jiang R, Li L, Wang C, Nussenzveig DR, et al (2019) Nucleoporin Seh1 Interacts with Olig2/Brd7 to Promote Oligodendrocyte Differentiation and Myelination. Neuron 102: 587–601.e7

Love MI, Huber W & Anders S (2014) Moderated estimation of fold change and dispersion for RNA-seq data with DESeq2. Genome Biol 15: 1–21

MacArthur IC, Bei Y, Garcia HD, Ortiz M V., Toedling J, Klironomos F, Rolff J, Eggert A, Schulte JH, Kentsis A, et al (2019) Prohibitin promotes dedifferentiation and is a potential therapeutic target in neuroblastoma. JCI Insight 4: e127130

Mamber C, Kamphuis W, Haring NL, Peprah N, Middeldorp J & Hol EM (2012) GFAPδ Expression in Glia of the Developmental and Adolescent Mouse Brain. PLoS One 7: 1–15

Markaki Y, Gan Chong J, Wang Y, Jacobson EC, Luong C, Tan SYX, Jachowicz JW, Strehle M, Maestrini D, Banerjee AK, et al (2021) Xist nucleates local protein gradients to propagate silencing across the X chromosome. Cell 184: 6174–6192.e32

Mase S, Shitamukai A, Wu Q, Morimoto M, Gridley T & Matsuzaki F (2021) Notch1 and Notch2 collaboratively maintain radial glial cells in mouse neurogenesis. Neurosci Res 170: 122–132

Mercer TR, Qureshi I a, Gokhan S, Dinger ME, Li G, Mattick JS & Mehler MF (2010) Long noncoding RNAs in neuronal-glial fate specification and oligodendrocyte lineage maturation. BMC Neurosci 11: 14

Mi H, Muruganujan A & Thomas PD (2013) PANTHER in 2013: Modeling the evolution of gene function, and other gene attributes, in the context of phylogenetic trees. Nucleic Acids Res 41: 377–386

Moffat J, Grueneberg D a., Yang X, Kim SY, Kloepfer AM, Hinkle G, Piqani B, Eisenhaure TM, Luo B, Grenier JK, et al (2006) A Lentiviral RNAi Library for Human and Mouse Genes Applied to an Arrayed Viral High-Content Screen. Cell 124: 1283–1298

Mukherjee S, Brulet R, Zhang L & Hsieh J (2016) REST regulation of gene networks in adult neural stem cells. Nat Commun 7: 1–14

Mukhtar T, Breda J, Grison A, Karimaddini Z, Grobecker P, Iber D, Beisel C, van Nimwegen E & Taylor V (2020) Tead transcription factors differentially regulate cortical development. Sci Rep 10: 1–19

Ng SY, Bogu GK, Soh BS & Stanton LW (2013) The long noncoding RNA RMST interacts with SOX2 to regulate neurogenesis. Mol Cell 51: 349–359

Palazzo AF & Koonin E V. (2020) Functional Long Non-coding RNAs Evolve from Junk Transcripts. Cell 183: 1151–1161

Pascual-Garcia P & Capelson M (2021) The nuclear pore complex and the genome: organizing and regulatory principles. Curr Opin Genet Dev 67: 142–150

Pascual-Garcia P, Debo B, Aleman JR, Talamas JA, Lan Y, Nguyen NH, Won KJ & Capelson M (2017) Metazoan Nuclear Pores Provide a Scaffold for Poised Genes and Mediate Induced Enhancer-Promoter Contacts. Mol Cell 66: 63–76.e6

Perez-Riverol Y, Bai J, Bandla C, García-Seisdedos D, Hewapathirana S, Kamatchinathan S, Kundu DJ, Prakash A, Frericks-Zipper A, Eisenacher M, et al (2022) The PRIDE database resources in 2022: a hub for mass spectrometry-based proteomics evidences. Nucleic Acids Res 50: D543– D552

Ponjavic J, Oliver PL, Lunter G & Ponting CP (2009) Genomic and Transcriptional Co-Localization of Protein-Coding and Long Non-Coding RNA Pairs in the Developing Brain. PLOS Genet 5: e1000617

Price RL, Bhan A & Mandal SS (2021) HOTAIR beyond repression: In protein degradation, inflammation, DNA damage response, and cell signaling. DNA Repair (Amst) 105: 103141

Quinlan AR & Hall IM (2010) BEDTools: A flexible suite of utilities for comparing genomic features. Bioinformatics 26: 841–842

Quinn JJ & Chang HY (2016) Unique features of long non-coding RNA biogenesis and function. Nat Rev Genet 17: 47–62

Raducu M, Fung E, Serres S, Infante P, Barberis A, Fischer R, Bristow C, Thézénas M, Finta C, Christianson JC, et al (2016) SCF (Fbxl17) ubiquitylation of Sufu regulates Hedgehog signaling and medulloblastoma development . EMBO J 35: 1400–1416

Rajalingam K & Rudel T (2005) Ras-Raf signaling needs prohibitin. Cell Cycle 4: 1503–1505

Ramírez F, Ryan DP, Grüning B, Bhardwaj V, Kilpert F, Richter AS, Heyne S, Dündar F & Manke T (2016) deepTools2: a next generation web server for deep-sequencing data analysis. Nucleic Acids Res 44: W160–W165

Ramos AD, Andersen RE, Liu SJ, Nowakowski TJ, Hong SJ, Gertz CC, Salinas RD, Zarabi H, Kriegstein AR & Lim DA (2015) The long noncoding RNA Pnky regulates neuronal differentiation of embryonic and postnatal neural stem cells. Cell Stem Cell 16: 439–447

Rani N, Nowakowski TJ, Zhou H, Godshalk SE, Lisi V, Kriegstein AR & Kosik KS (2016) A Primate lncRNA Mediates Notch Signaling during Neuronal Development by Sequestering miRNA. Neuron 90: 1174–1188

Rinn JL & Chang HY (2012) Genome regulation by long noncoding RNAs. Annu Rev Biochem 81: 145–166

Rinn JL & Chang HY (2020) Long Noncoding RNAs: Molecular Modalities to Organismal Functions. Annu Rev Biochem 89: 283–308

Rutenberg-Schoenberg M, Sexton AN & Simon MD (2016) The Properties of Long Noncoding RNAs That Regulate Chromatin. Annu Rev Genomics Hum Genet 17: 69–94

Sauer B (1994) Site-specific recombination: developments and applications. Curr Opin Biotechnol 5: 521–527

Schoenherr CJ & Anderson DJ (1995) The neuron-restrictive silencer factor (NRSF): a coordinate repressor of multiple neuron-specific genes. Science 267: 1360–1363

See RH, Caday-Malcolm RA, Singaraja RR, Zhou S, Silverston A, Huber MT, Moran J, James ER, Janoo R, Savill JM, et al (2002) Protein kinase A site-specific phosphorylation regulates ATP-binding cassette A1 (ABCA1)-mediated phospholipid efflux. J Biol Chem 277: 41835–41842

Shin E, Kashiwagi Y, Kuriu T, Iwasaki H, Tanaka T, Koizumi H, Gleeson JG & Okabe S (2013) Doublecortin-like kinase enhances dendritic remodelling and negatively regulates synapse maturation. Nat Commun 4: 1440

Statello L, Guo CJ, Chen LL & Huarte M (2021) Gene regulation by long non-coding RNAs and its biological functions. Nat Rev Mol Cell Biol 22: 96–118

van Steensel B & Belmont AS (2017) Lamina-Associated Domains: Links with Chromosome Architecture, Heterochromatin, and Gene Repression. Cell 169: 780–791

Sueda R & Kageyama R (2019) Regulation of active and quiescent somatic stem cells by Notch signaling. Dev Growth Differ 62: 59–66

Takagi M, Sueishi M, Saiwaki T, Kametaka A & Yoneda Y (2001) A Novel Nucleolar Protein, NIFK, Interacts with the Forkhead Associated Domain of Ki-67 Antigen in Mitosis. J Biol Chem 276: 25386–25391

Tsai BP, Wang X, Huang L & Waterman ML (2011) Quantitative profiling of in vivo-assembled RNA-protein complexes using a novel integrated proteomic approach. Mol Cell Proteomics 10: M110.007385

Tusher VG, Tibshirani R & Chu G (2001) Significance analysis of microarrays applied to the ionizing radiation response. Proc Natl Acad Sci U S A 98: 5116–5121

Tyanova S, Temu T, Sinitcyn P, Carlson A, Hein MY, Geiger T, Mann M & Cox J (2016) The Perseus computational platform for comprehensive analysis of (prote)omics data. Nat Methods 13: 731– 40

Vertii A, Ou J, Yu J, Yan A, Pagès H, Liu H, Zhu LJ & Kaufman PD (2019) Two contrasting classes of nucleolus-associated domains in mouse fibroblast heterochromatin. Genome Res 29: 1235– 1249

Wang J, He X, Luo Y & Yarbrough WG (2006) A novel ARF-binding protein (LZAP) alters ARF regulation of HDM2. Biochem J 393: 489–501

Wang L, Hou S & Han Y-G (2016) Hedgehog signaling promotes basal progenitor expansion and the growth and folding of the neocortex. Nat Neurosci 19: 888–896

Wang S, Fusaro G, Padmanabhan J & Chellappan SP (2002) Prohibitin co-localizes with Rb in the nucleus and recruits N-CoR and HDAC1 for transcriptional repression. Oncogene 21: 8388–8396

Wang T, Ma X, Zhu H, Li A, Du G & Chen J (2012) Available methods for assembling expression cassettes for synthetic biology. Appl Microbiol Biotechnol 93: 1853–1863

Werner MS & Ruthenburg AJ (2015) Nuclear Fractionation Reveals Thousands of Chromatin-Tethered Noncoding RNAs Adjacent to Active Genes. Cell Rep 12: 1089–1098

Wiśniewski JR, Zougman A, Nagaraj N & Mann M (2009) Universal sample preparation method for proteome analysis. Nat Methods 6: 359–362

Xi J, Xu Y, Guo Z, Li J, Wu Y, Sun Q, Wang Y, Chen M, Zhu S, Bian S, et al (2022) LncRNA SOX1- OT V1 acts as a decoy of HDAC10 to promote SOX1-dependent hESC neuronal differentiation . EMBO Rep 23: 1–17

Yao PJ, Petralia RS & Mattson MP (2016) Sonic Hedgehog Signaling and Hippocampal Neuroplasticity. Trends Neurosci 39: 840–850

Yao RW, Wang Y & Chen LL (2019) Cellular functions of long noncoding RNAs. Nat Cell Biol 21: 542– 551

Yuan B, Latek R, Hossbach M, Tuschl T & Lewitter F (2004) siRNA selection server: An automated siRNA oligonucleotide predicition server. Nucleic Acids Res 32: 130–134

Zhou Z, Licklider LJ, Gygi SP & Reed R (2002) Comprehensive proteomic analysis of the human spliceosome. Nature 419: 182–185

Zhou Z & Reed R (2003) Purification of Functional RNA-Protein Complexes using MS2-MBP. Curr Protoc Mol Biol 63: 27.3.1-27.3.7

Ziller MJ, Edri R, Yaffe Y, Donaghey J, Pop R, Mallard W, Issner R, Gifford CA, Goren A, Xing J, et al (2015) Dissecting neural differentiation regulatory networks through epigenetic footprinting. Nature 518: 355–359

Zimmer-Bensch G (2019) Emerging Roles of Long Non-Coding RNAs as Drivers of Brain Evolution. Cells 8

